# Reclassifying Cancer: Defining tumour cell cycle activity in terms of its tissue of origin in over 13,000 samples

**DOI:** 10.1101/2022.02.15.480623

**Authors:** Arian Lundberg, Joan Jong Jing Yi, Linda S. Lindström, Nicholas P. Tobin

**Affiliations:** Department of Oncology and Pathology, Karolinska Institutet and University Hospital, Stockholm, Sweden; Department of Radiation Oncology, University of California - San Francisco, San Francisco, CA, USA; Department of Radiation Oncology, Stanford School of Medicine, Stanford, CA, USA; School of Biological Sciences, Nanyang Technological University, Singapore

**Keywords:** Cell cycle, pan-cancer, gynaecological, normal tissue, genomic analyses

## Abstract

Genomic alterations resulting in loss of control over the cell cycle is a fundamental hallmark of human malignancies. Whilst pan-cancer studies have broadly assessed tumour genomics and their impact on oncogenic pathways, analyses taking the baseline signalling levels in normal tissue into account are lacking. To this end, we aimed to reclassify the cell cycle activity of tumours in terms of their tissue of origin and determine the DNA mutations, chromosome arm-level changes and signalling pathways driving cell cycle activity. Combining normal tissue and pan-cancer data from over 13,000 samples we demonstrate that tumours of gynaecological origin show the highest levels of baseline corrected cell cycle activity, partially owing to hormonal signalling and gene expression changes. We also show that normal and tumour tissues can be separated into groups (quadrants) of low/high cell cycle activity and propose the novel hypothesis of an upper limit on these activity levels in tumours.

## Introduction

The cell cycle is an all-encompassing term that describes the way by which a cell duplicates its genetic contents and subsequently divides into two identical daughter cells. The cycle consists of two main events – interphase and the mitotic or M phase(1). Interphase can be subdivided into the S phase, where DNA replication occurs, and either side of this come the Gap 1 (G1) and 2 (G2) phases. In addition to the G1, S, G2 and M phases the cell can also enter a quiescent or non-proliferative state after prolonged serum withdrawal termed G0(2). Reintroduction of serum allows the cell to again enter the cell cycle at G1. Transitions between the cell cycle phases are governed by the cyclin-dependent kinases (CDKs) and their binding partners from the cyclin family of proteins, cyclin D, E, A and B.

Loss of control over the cell cycle resulting in uncontrolled proliferation is a hallmark of cancer(3) and aberrant expression of cell cycle-related genes is frequently observed at a pan-cancer level(4). The most common cell cycle gene alterations include deletion of the p16 tumour suppressor gene (*CDKN2A/B*, deleted in approximately 20% of all cancer patients), deletions and mutations in the retinoblastoma protein (*RB1*, 7% of patients) and amplifications of the cyclin D1 (*CCND1*, 6% of all cancer patients, up to 30% in breast cancer(5)), E1 (*CCNE1*, 3.6%) and *CDK4* (3.2%) genes(6). In line with this, pan-cancer studies have also indicated that genomic aberrations are more common in signalling pathways that promote S phase entry or prevent cell cycle exit (e.g. DNA damage response pathway)(4) rather than those associated with mitotic entry and exit (for review see here(1)).

We recently conducted a comprehensive pan-cancer analysis of the genomic aberrations present in tumour samples based on the magnitude of cell cycle pathway activity(7). We found that cell cycle activity varies broadly across and within the cancer types of The Cancer Genome Atlas (TCGA) pan-cancer cohort and that *TP53, PIK3CA* and chromosomal alterations occur with increasing frequency in tumours with increasing cell cycle activity. Here, we build on this work, hypothesising that the starting or baseline level of cell cycle activity in normal tissue influences the level of tumour cell cycle activity and by extension, that we can reclassify tumour cell cycle activity by placing it in terms of its normal tissue of origin. To test this hypothesis, we analyse over 13,000 samples from the UCSC Toil RNA-seq Recompute Compendium of batch corrected RNA-seq data from the Genotype-Tissue Expression (GTEx, normal tissue) and pan-cancer TCGA projects(8). Specifically, we apply a gene expression signature representative of the cell cycle (the Cell Cycle Score, CCS(7,9,10)) to the samples from the GTEx and TCGA studies and recalculate the cell cycle activity of 23 different cancer types taking normal tissue baseline cell cycle levels into account. We also compare the two groups of cancers with the lowest and highest baseline corrected cell cycle activity in order to understand the specific signalling pathways and genomic aberrations driving this activity difference.

## Materials and Methods

### Study population and specimens

The UCSC Toil RNA-seq Recompute Compendium is a collection of batch effect-corrected RNA-seq samples from three datasets including The Cancer Genome Atlas (TCGA) (N = 10535), Therapeutically Applicable Research To Generate Effective Treatments (TARGET) (N = 734) and Genotype-Tissue Expression (GTex) (N = 7862)(8). The compendium contains tumours from 24 different cancer types, including Adrenocortical carcinoma (ACC), Bladder Urothelial Carcinoma (BLCA), Brain lower grade Glioma (LGG), Breast invasive carcinoma (BRCA), Cervical squamous cell carcinoma and endocervical adenocarcinoma (CESC), Colon adenocarcinoma (COAD), Esophageal carcinoma (ESCA), Glioblatoma multiforme (GBM), Head and Neck squamous cell carcinoma (HNSC), Kidney Chromophobe (KICH), Kidney renal clear cell carcinoma (KIRC), Kidney renal papillary cell carcinoma (KIRP), Liver hepatocellular carcinoma (LICH), Lung adenocarcinoma (LUAD), Lung squamous cell carcinoma (LUSC), Ovarian serous cystadenocarcinoma (OV), Pancreatic adenocarcinoma (PAAD), Prostate adenocarcinoma (PRAD), Skin Cutaneous Melanoma (SKCM), Stomach adenocarcinoma (STAD), Testicular Germ Cell tumours (TGCT), Thyroid carcinoma (THCA), Uterine Carcinosarcoma (UCS), and Uterine Corpus Endometrial Carcinoma (UCEC) as well as normal tissue samples. All data are publicly available(11) and the quality control, normalization and gene level counts were performed by the Toil investigators as described in the original publication(8).

### Cell Cycle Score (CCS) and Baseline Corrected-Cell Cycle Score (BC-CCS)

The CCS signature, along with its gene composition and method of application, has been previously extensively described(7,9,10). Briefly, the signature is comprised of 463 cell cycle-related genes that were originally identified through the aggregation of three different pathway-related databases – Kyoto Encyclopedia of Genes and Genomes (KEGG), HUGO gene nomenclature committee (HGNC) and Cyclebase 3.0(12–14). 446 of the 463 original CCS signature genes were present in this study (Supplemental Table 1) and the final signature score was derived by summing up expression values of the signature genes resulting in a single CCS value on a per tissue sample/ tumour basis. As these samples have already been normalised and standardised together as part of the Toil pipeline to make them directly comparable, the Baseline Corrected - Cell Cycle Score (BC-CCS) was calculated for each tumour sample by subtracting the median CCS of its GTEx normal tissue from the CCS of the tumour sample. For example: *BC-CCS Bladder Tumour 1 = CCS Bladder Tumour 1 - median CCS all normal Bladder Tissue.* The BC-CCS was additionally adjusted for tumour purity to account for differences in tumour epithelial content using the ABSOLUTE algorithm values previously derived by the pan-cancer investigators(15) and shown in Supplemental Table 2. Finally, both the CCS and BC-CCS continuous variables were scaled to values between 0 and 1 to aid with plotting and data visualisation.

### Batch effect and outlier assessment

Principal Component Analysis (PCA) was performed with the *gmodels* R-package using (i) the genes with the highest variation in the dataset (N = 6364) and (ii) the genes included in the Cell Cycle Score (CCS, N = 446)(9). Batch effects and the presence of outliers were assessed through manual examination of the PCA plots and overlaying study origin (GTEx, TCGA-normal, TCGA-cancer), tissue type (each of the 19 tissue types included in the study) or baseline corrected change in Cell Cycle Score data. Uniform Manifold Approximation and Projection (UMAP) was also applied to the dataset using the *umap* R-package as it provides high accuracy in separating the features of a complex data while identifying batch effects(16).

### Mutation and chromosomal arm-level analyses

We used publicly available fully processed(17) mutation and chromosomal arm-level alteration data from the TCGA pan-cancer samples housed in the Genome Data Commons (GDC) database (https://gdc.cancer.gov). These data were used to compare the DNA-level differences between the two groups with the lowest and highest BC-CCS, adjusting for the number of individual tumours with each cancer type of these two groups (Group 1: HNSC, KICH, KIRP UCEC and Group 2: CESC, OV, UCS) and for multiple testing using False Discovery Rate (FDR). Of note, for the mutational analysis we focused on 299 cancer driver genes, that were manually annotated by experts in the field(17).

### Gene expression and Gene Set Enrichment analyses

Differential gene expression analysis was performed using the *limma*(18) R-package in order to understand the mRNA differences between the two groups with the lowest and highest BC-CCS. The model matrix included additional variables to adjust for individual cancer types (HNSC, KICH, KIRP UCEC, CESC, OV, UCS), and the estrogen receptor gene *ESR1* as a continuous variable in order to adjust for the general impact of sex hormones on the cell cycle. Results were corrected for multiple testing and genes with Log_2_ fold-change (FC) > 2 and FDR < 5% were considered significant. Note that we applied this strict 5% FDR in order to derive a list of genes we could be more confident were differentially expressed. Gene Set Enrichment Analyses (GSEA) was used to evaluate the enrichment of cancer hallmark pathways within the differentially expressed genes using the *fgsea(19)* R-package. Pathway rankings were ordered based on FC and p values. Results were corrected for multiple testing and again we denote a stringent FDR < 10% threshold as statistical significance (GSEA uses 25% FDR as standard(20)).

### Statistical Analysis

Student’s t-test was used to assess differences between normal and tumour continuous CCS and Fisher-exact test was applied to determine if gene mutations or chromosomal arm-level alterations were significantly different between the two BC-CCS groups. Genes found to be differentially expressed at the mRNA level between the same groups were tested for their correlation to the BC-CCS using Spearman’s rank correlation test. All tests were 2-sided and *p* < 0.05 was considered as statistically significant. The data fulfilled the preconditions/ assumption of the above tests. Continuous CCS was normally distributed with low variation between groups. All statistical analyses were performed using R statistical software version 4.1.1(21).

## Results

### GTEx and TCGA samples cluster together on the basis of tissue of origin and cell cycle activity levels

In order to compare cell cycle activity between normal and cancerous tissues, we analysed the GTEx and TCGA pan-cancer datasets which were reprocessed, normalised and batch-corrected together as a part of the Toil recompute project(8). From the original 19,131 samples we filtered out those that were not a part of these two datasets (N = 734), that lacked mRNA-expression data (N = 92), that did not have representative tissue from the same site in both studies (N = 802) or where the normal tissue or cancer site contained fewer than ten samples (N = 4,043, Figure 1). Next, we assessed possible batch effects and the presence of outliers with PCA plots using RNA-seq data from the remaining 13,460 samples. Plotting on the basis of the most variable genes across all samples, PCA showed a long tail of samples stretching into the lower right quadrant (Supplemental Figure 1A coloured in red, and Supplemental Figure 1B). Closer inspection and annotation of these samples showed all were GTEx normal testicular tissue (without the presence of TCGA testicular tumours), indicating a potential batch effect and as such all normal and cancer testicular samples were removed from further analyses (N = 319). Similarly, after applying our Cell Cycle Score (CCS) signature to the remaining samples we noted a small number of outliers (N = 24) from mixed origin using PCA (Supplemental Figure 1C and D, circled in red) which were also removed from further analysis leaving 13,117 samples in total. A breakdown of the origin and number of all removed samples is shown in Supplemental Table 3.

**Figure 1.**
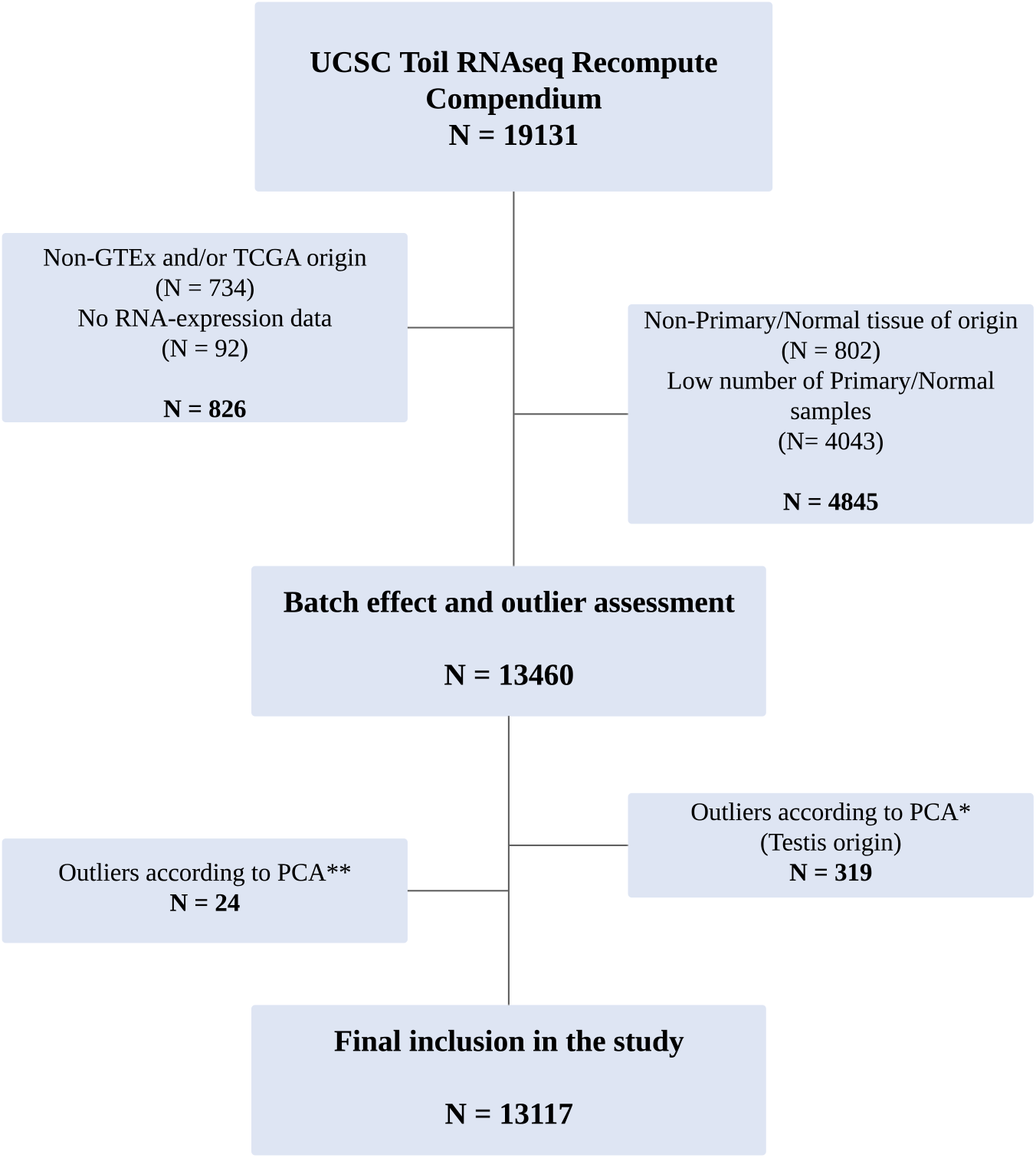
CONSORT diagram of sample filtering and selection. *Excluded samples as shown in Supplemental figure 1A/B; **excluded samples as shown in Supplemental figure 1C/D.

Replotting the PCA of most varying genes in 13,117 samples (N = 4,979 normals and 8,138 cancers) did not show any clear grouping on the basis of study origin (Figure 2A; grey = GTEx, light orange = TCGA normal tissue and blue = TCGA tumours) but rather on the basis of tissue type (Figure 2B). A similar finding was observed using UMAP (Supplemental Figure 1E and F). These results indicate an absence of additional batch effects and support further direct comparison of tissue types. Using the CCS, samples clustered into large groups on the basis of GTEx and TCGA normal samples or TCGA tumour samples (Figure 2C, compare grey and light orange colours vs. blue), whilst tissue types generally stayed in close proximity (Figure 2D). This result was anticipated when plotting based on cell cycle genes as tumours will in general cycle more quickly and demonstrate higher proliferation levels. Similar results were found using the CCS genes and UMAP (Figure 2E and F) and a core group of samples high in CCS was also readily apparent (Figure 2G, higher CCS = yellow). Together, these results indicate that GTEx and TCGA samples cluster together on the basis of tissue of origin (as expected, given the aim of the Toil recompute project) and also group on the basis of their cell cycle activity levels.

**Figure 2.**
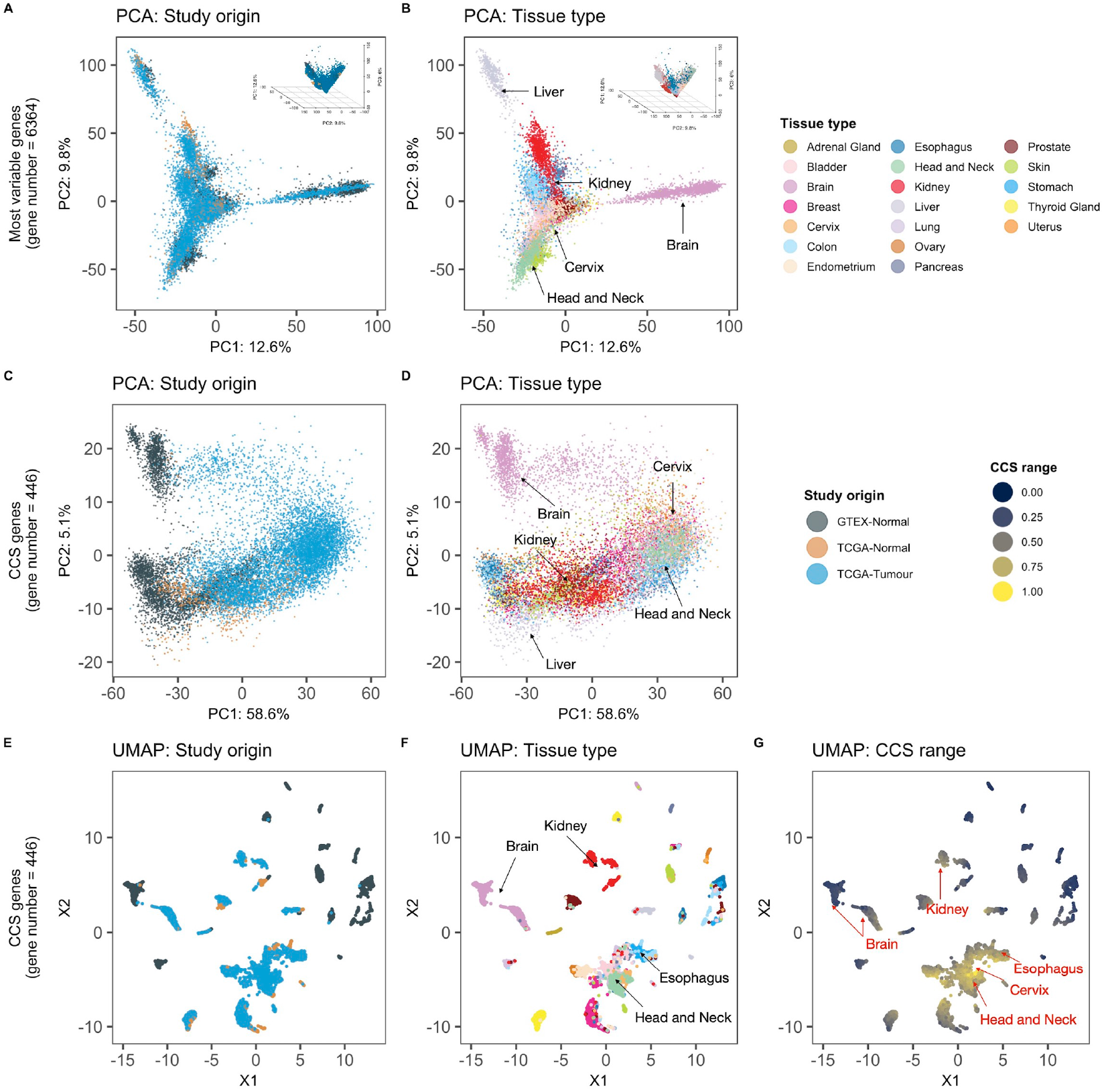
PCA and UMAP dimensionality reduction of GTEx and Pan-Cancer Atlas datasets. Toil pipeline batch-corrected mRNA expression data from the GTEx and TCGA pan-cancer datasets (N = 13117 in total) were used to perform A) Principle component analyses (PCA) using the most variable genes in the dataset with an overlay of colours based on study origin or B) Tissue type; C) PCA using the genes of the CCS signature with an overlay of colours based on study origin or D) Tissue type; E) Uniform Manifold Approximation and Projection (UMAP) using the genes of the CCS signature with an overlay of colours based on study origin or F) Tissue type or G) CCS range.

### Tumours from gynaecological tissues show the highest baseline corrected cell cycle activity levels

To visualise the differences more clearly in cell cycle activity between tumours and their normal tissue of origin, we plotted the CCS as a continuous variable using boxplots and separated samples into tissue of origin. While normal samples showed varying CCS levels when compared to each other (Figure 3A and Supplemental Figure 2A), tumour samples had a consistently higher CCS relative to their tissue of origin (Figure 3A, P < 0.001, Students’ T-test, adjusted for multiple testing, first boxplot in each tissue type is normal tissue and the rest are tumours). The importance of relating tumour cell cycle activity to normal tissue is also evident from these boxplots: Bladder cancer (“BLCA”, red arrow) has higher cell cycle activity than glioblastoma multiforme (“GBM”, blue arrow), but normal levels of bladder cell cycle activity are much higher than that of brain tissue (compare “Bladder” to “Brain”, black arrows). This means that at the absolute level BLCA shows higher cell cycle activity levels than GBM, but at a relative level (relative to its baseline level in normal tissue) GBM’s cell cycle activity is much higher than BLCA. Next, we extended this concept by reclassifying all tumour samples taking (subtracting) the baseline cell cycle activity of its normal tissue of origin into account to give a new Baseline Corrected-Cell Cycle Score (BC-CCS). As the tumour epithelial cell content differs for different cancer types (Supplemental Table 2), we also adjusted the BC-CCS for tumour purity using values derived from the ABSOLUTE algorithm. Notably, we found that in general, cancers of gynaecological origin (Cervical squamous cell carcinoma and endocervical adenocarcinoma, CESC; Ovarian serous cystadenocarcinoma, OV; and Uterine Carcinosarcoma, UCS) display the highest BC-CCS levels and Head and Neck squamous cell carcinoma (HNSC), Kidney Chromophobe (KICH) and Kidney renal papillary cell carcinoma (KIRP) the lowest (Figure 3B). This finding is even more interesting when placed in the context of our previous research in the TCGA pan-cancer atlas tumours only without baseline correction, where we found HNSC to be a tumour type with one of the highest CCS activity levels(7). We show here that head and neck tissue also has the highest level of cell cycle activity in normal tissue (Supplemental Figure 2A). Taken together, this may imply that cell cycle activity in head and neck cancers simply cannot be pushed much higher as its baseline levels are already so high in normal tissue that it has reached its upper limit or “ceiling”. Conversely, gynaecological cancers start at a much lower baseline level (Supplemental Figure 2A, see Uterus, Ovary, Cervix), giving them a higher ceiling to proliferate into once an oncogenic event has occurred. The relationship between cell cycle activity in normal and tumour tissues based on median CCS expression is visualized in a scatterplot divided into quadrants of high/low CCS in Figure 3C. Here, bladder, endometrial and head and neck tissues all show high normal/ high tumour cell cycle activity, whereas cervix, oesophagus, uterus and ovary tissue show low normal/ high tumour cell cycle activity. The BC-CCS is also visualised anatomically in Supplemental Figure 3 and similar pan-cancer boxplots where breast cancer is separated into its molecular subtypes are also shown in Supplemental Figure 2B and C.

**Figure 3.**
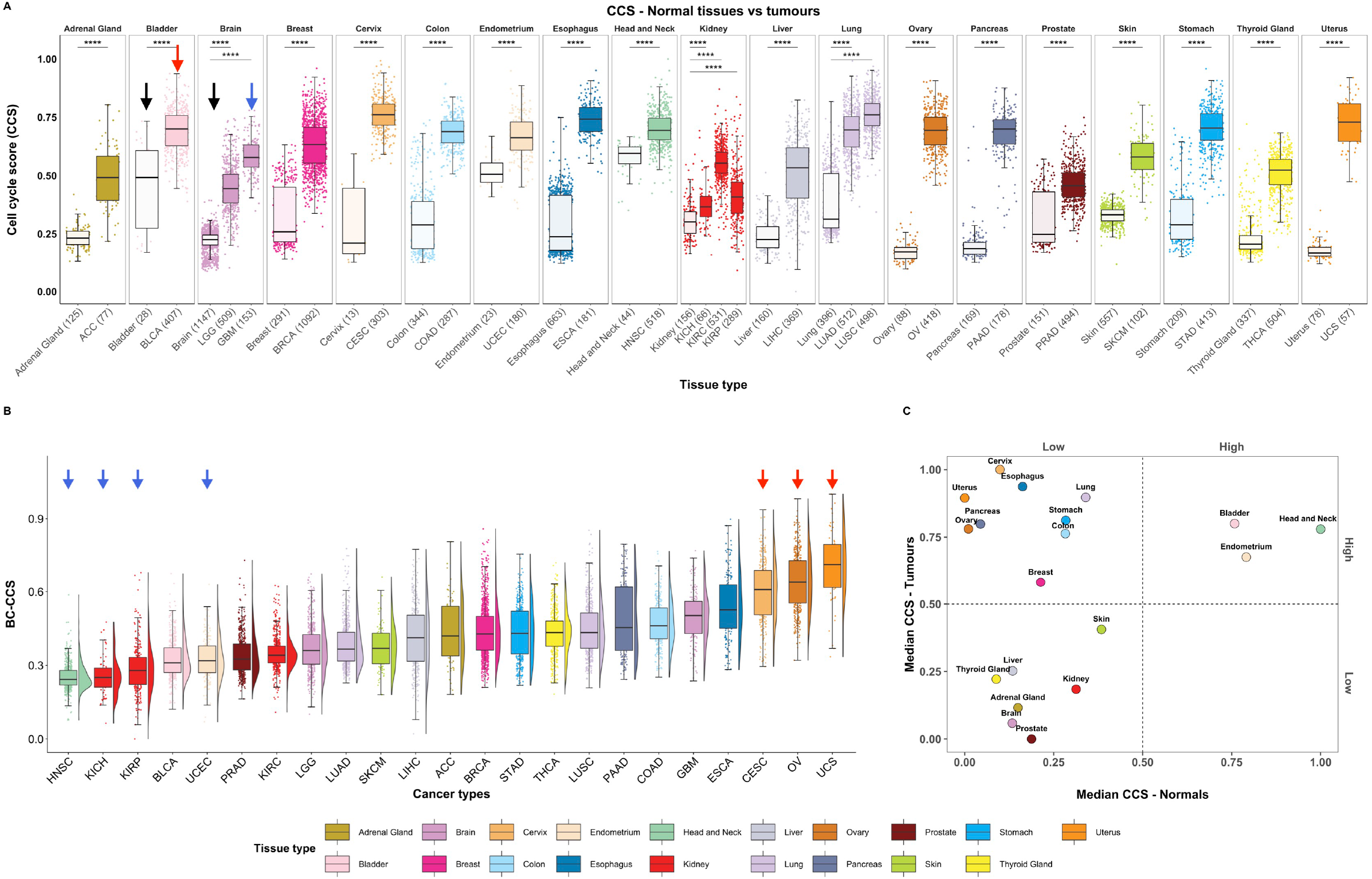
Boxplots and scatterplot comparing cell cycle activity across normal tissue and Pan-Cancer Atlas tumour samples. Toil pipeline batch corrected mRNA expression data from the GTEx and TCGA pan-cancer datasets were used to examine cell cycle activity. A) The CCS gene expression signature was used to create boxplots within in each normal and cancer tissue type, black arrows denote normal bladder and brain tissue, red= BLCA, blue = GBM. B) The baseline corrected-CCS (BC-CCS) in pan-cancer tumours, where the median CCS value for the normal tissue of origin has been subtracted from each cancer type. Density plot is placed beside each boxplot. Blue and red arrows indicate cancers from Groups 1 and 2, respectively. C) Scatterplot showing the relationship of cell cycle activity in normal tissues with tumour samples on the basis of median CCS expression. Median CCS expression standardized by scaling between 0-1. *p* values in boxplots are based on Student’s T-test and corrected for multiple testing; **** = *p* < 0.001.

### Statistically significant differences in DNA copy number when comparing the highest and lowest BC-CCS tumour groups

Next, we wanted to understand if specific signalling pathways or genomic aberrations are more common in tumour types showing a higher BC-CCS compared to those showing lower. For this, we selected the tumour types at the extremes of Figure 3B, forming two comparison groups (Figure 3B, red and blue arrows). Group 1 consisted of tumour types showing the lowest BC-CCS (HNSC, KICH and KIRP) with the addition of Uterine Corpus Endometrial Carcinoma (UCEC). UCEC is an exception to our findings in Figure 3B in that it is a gynaecological tumour type that shows a low BC-CCS. Given that the three highest BC-CCS tumour groups that form Group 2 (CESC, OV, UCS) are all of gynaecological origin, we wanted to make sure that any potential differences found when comparing Group 1 to Group 2 were not only due to a comparison of gynaecological vs. non-gynaecological tissues. For this reason, we included UCEC in Group 1 as an internal control.

First, we focused on a comparison of DNA-level mutations in 299 known cancer driver genes(17) between these two groups. The top 20 mutations after adjusting for number in each group are shown in Figure 4A and as anticipated given their high pan-cancer mutation frequency, the *TP53, PIK3CA* and *PTEN* genes are amongst the most mutated in both groups. Using a Fisher-exact test to assess whether any mutations were statistically more common in Group 2, we found that majority of differences at an individual gene level occurred in a low percentage of tumours (< 5%) within each cancer type (Figure 4C). One exception to this was the *FBXW7* gene which was mutated in 54% of tumours in Group 2 vs. 31% in Group 1, however this result is mostly driven by the UCS in Group 2 where 40% of tumour carried this mutation. Second, and continuing our DNA-level analysis, we assessed chromosomal arm-level copy number differences between the same groups. After adjusting for tumour numbers within each cancer type, we found increased deletions in 16 different chromosomal arms in Group 2 relative to Group 1 (Figure 4D, Fisher-exact test, FDR < 5% for all comparisons). The largest differences in deletions were found in arms 16q, 8p, 9q, 22q and 4q (Figure 4D) and similarly, of the 11 different chromosomal arms found to have significantly increased amplifications between the two groups, the largest differences were found in 3q, 1q, 5p, 6p and 2p (Figure 4E, Fisher-exact test and FDR < 5% for all comparisons). However, semi-supervised clustering on the basis of these amplifications and deletions failed to cluster the tumours of Groups 1 and 2 or all other pan-caner samples into subgroups with lower or higher BC-CCS (Supplemental Figure 4). This implies that these arm-level genomic alterations are unlikely to drive the differences in baseline corrected tumour cell cycle activity at a pan-cancer level.

**Figure 4.**
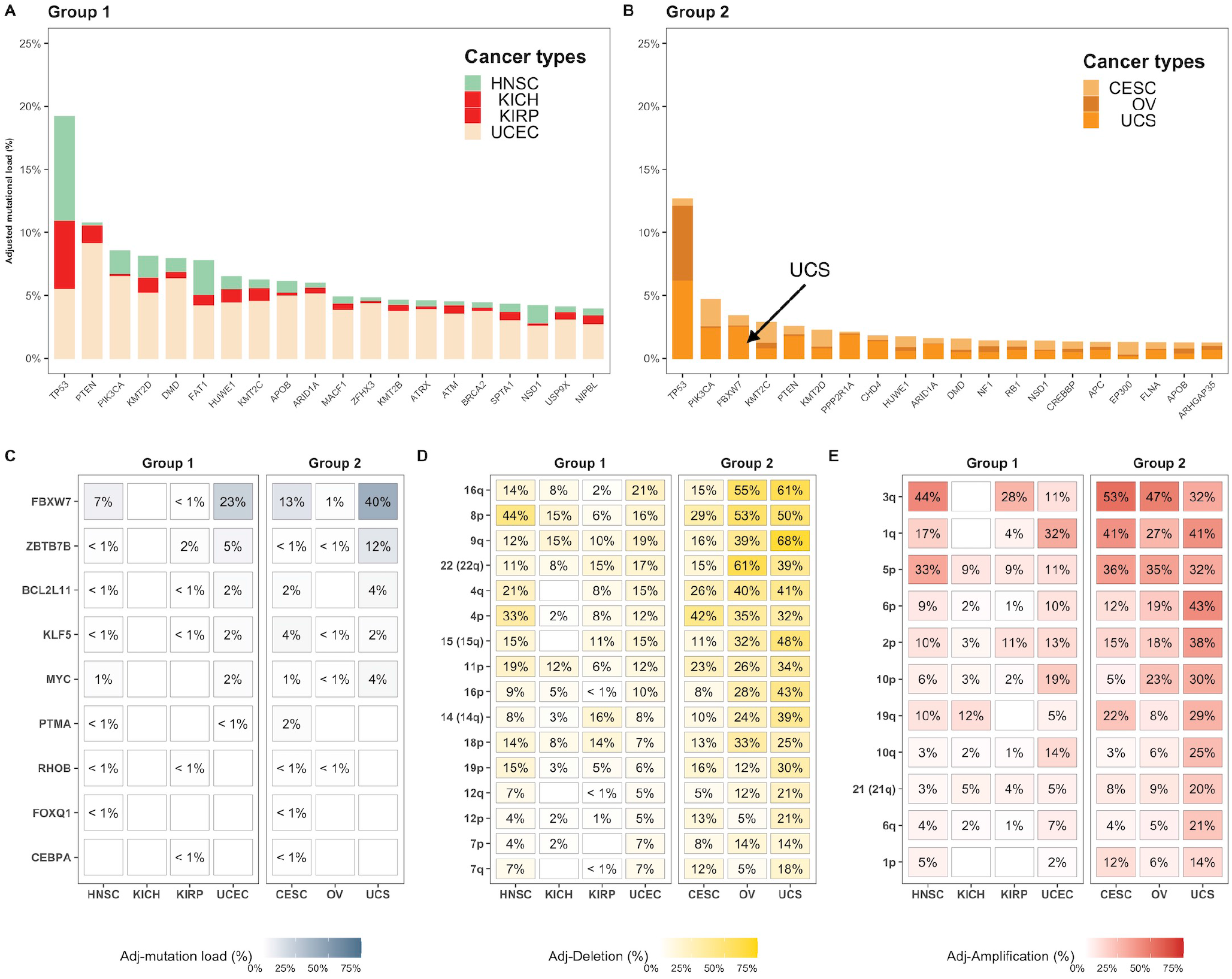
Most highly mutated genes and chromosomal arm-level aberrations in Groups 1 and 2. The cancer types with the highest and lowest BC-CCS were placed into two groups: Group 1 = HNSC, KICH, KIRP and UCEC; Group 2 = CESC, OV and UCS. A) The top 20 most mutated cancer driver genes for Group 1 and B) Group 2, mutation percentage has been adjusted for sample number within each cancer type for all genes individually C) Driver genes with a higher mutation level in Group 2 relative to Group 1 by Fisher’s Exact Test. D) Chromosomal arm-level deletions that occur with higher frequency in Group 2 relative to Group 1 and E) Chromosomal arm-level amplifications occurring with higher frequency in Group 2 relative to Group 1. All comparisons were adjusted for multiple testing.

### Statistically significant differences in hormone signalling and gene expression when comparing the highest and lowest BC-CCS tumour groups

As the sex hormones are known mitogens with an established role in pan-gyn studies(22) and in driving the cell cycle, we next determined whether the mRNA levels of the estrogren receptor alpha (ER-α), progesterone (PR) or androgen receptor (AR) genes were different between Group 1 and Group 2. We found that genes and gene modules (groups of genes) representative of all three hormones show higher levels in Group 2 (Figure 5A, Students T-test). When considering all pan-cancer samples, however, the correlation between the three gene modules and the BC-CCS was very weak (see inset of Supplemental Figure 5, R = 0.047, 0.095 and 0.083 for the estrogen, progesterone and androgen gene modules respective, Spearman’s correlation). In addition, boxplots of the same gene modules in individual cancer types show that not all cancer types with a high BC-CCS have high expression of sex hormones (Supplemental Figure 5A-C). These findings imply that while sex hormone expression may partially explain the increase in BC-CCS in some tumour types, it does not account for a high BC-CCS across all cancers. This is to be expected as sex hormone exposure has not been shown to be a risk factor for all cancer types.

**Figure 5.**
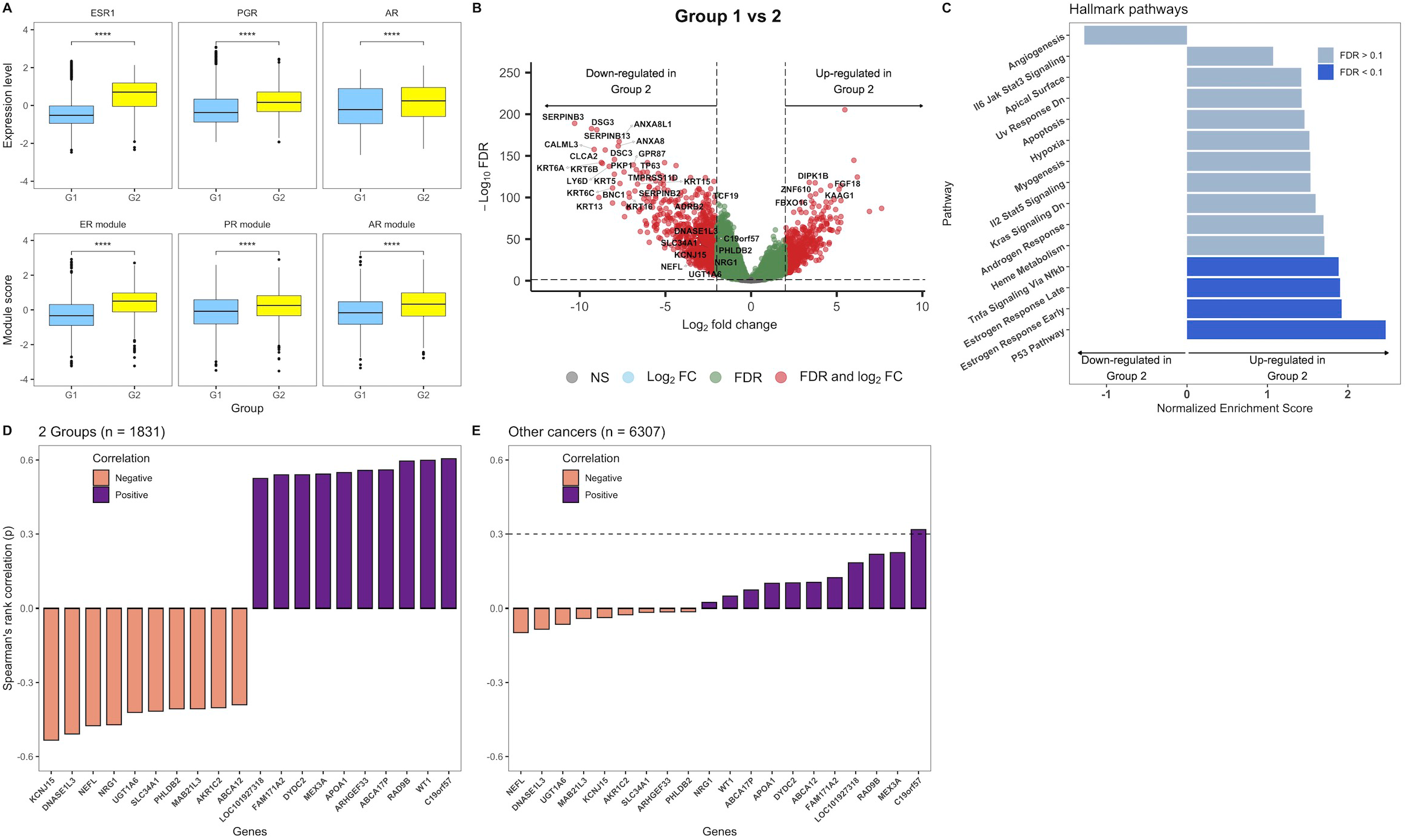
Comparison of sex hormone gene modules, differentially expressed genes and signalling pathways between Groups 1 and 2. mRNA data from the tumours of Groups 1 and 2 was used to compare: A) Expression of ER-α, Progesterone and Androgen receptor genes and representative gene modules B) Differential gene expression between the two groups, a volcano plot representing genes that are up- or down-regulated in Group 2 relative to Group 1 is shown C) Gene set enrichment analyses (GSEA) on the basis of the differentially expressed genes in Group 2. D) Barplots showing the genes with the highest Spearman’s rank correlation to the baseline corrected-Cell Cycle Score (BC-CCS) in the tumours of Groups 1 and 2 only (N = 1831) and E) Barplots showing the genes with the highest Spearman’s rank correlation to BC-CCS in all other cancer types of the pan-cancer cohort (minus the tumours from Groups 1 and 2, N= 6307). *p* values in boxplots are based on Student’s T-test and, were corrected for multiple testing; **** = *p* < 0.001. In the volcano plot genes with FDR < 5% and log_2_ fold-change > 2 or < −2 were considered significant. GSEA results with FDR < 10% were considered significant. Spearman’s rank correlation *Rho* was used for the correlation barplots; dashed line indicates *Rho* > 0.3.

To further examine the genes and pathways altered between Groups 1 and 2, we performed differential gene expression analysis, adjusting for both cancer type and the *ESR1* (ER-α) gene. The latter was included in an attempt to mitigate finding genes that were only representative of the sex hormone differences identified in Figure 5A. 522 genes were upregulated and 876 downregulated in Group 2 relative to Group 1 (Figure 5B). Pathway analysis of these genes showed upregulation of the p53, TNFα and, despite adjusting for the *ESR1* gene, estrogen response pathways (Figure 5C, Dark blue = significant at 10% FDR). We also examined individual correlations of the differentially expressed genes to the BC-CCS in all tumours of Groups 1 and Group 2 together to determine if any genes were driving the difference in BC-CCS. *C19orf57*, *WT1*, and *RAD9B* were the top 3 most positively and *KCNJ15*, *DNASE1L3* and *NEFL* the most negatively correlated to BC-CCS (Figure 5D, Spearman’s rank correlation). When correlating the same genes to BC-CCS in the rest of the pan-cancer cohort only *C19orf57* showed a moderate correlation (Figure 5D, Spearman’s *Rho* = 0.32)). Together, these results again point to a strong influence of estrogen signalling on the BC-CCS in gynaecological cancers and a potential role for the *C19orf57* gene in driving BC-CCS at a pan-cancer level.

## Discussion

In this study, we utilize RNA-seq data from two large public databases of normal and tumour tissue in order to reclassify cancer cell cycle activity in terms of its tissue of origin in over 13,000 samples. Following reclassification, we assessed the DNA mutation, chromosomal copy number and biological pathway signalling differences between tumours with the largest and smallest baseline corrected cell cycle changes, focusing on tumours of gynaecological origin. Our findings show first, that samples grouped broadly together on the basis of their tissue of origin or cell cycle activity level regardless of whether they originated from the GTEx or TCGA datasets. Second, that while normal samples showed varying CCS levels when compared to each other, tumour samples had a consistently higher CCS relative to their tissue of origin. Third, that in general, tumours of gynaecological origin (CESC, OV, UCS) show the highest baseline corrected cell cycle change. Fourth, that chromosomal arm-level alterations, hormone receptor signalling and specific genes including *C19orf57* are associated with this high cell cycle activity in gynaecological tumours and finally, that *C19orf57* also shows a moderate association with BC-CCS at a pan-cancer level.

We and others have previously shown that applying the CCS(7) or cell cycle-related classifiers(23) to the TCGA pan-cancer dataset separates tumours into those with high (Testicular Germ Cell tumours - TCGT, Head and Neck squamous cell carcinoma – HNSC and Cervical squamous cell carcinoma and endocervical adenocarcinoma - CESC) and low (Kidney Chromophobe – KICH, Kidney renal papillary cell carcinoma - KIRC, Thyroid carcinoma - THCA) cell cycle activity. Here, we demonstrate that by placing tumour cell cycle activity in terms of its normal tissue of origin, we gain a much clearer understanding of the genomic aberrations and biological signalling pathways that drive the largest changes in cell cycle activity. Indeed, it is striking that the gynaecological tissue types showing the lowest levels of cell cycle activity in normal tissue (OV, UCS - Supplemental Figure 2A) show the highest baseline corrected levels when examining their oncogenic counterpart. Conversely, head and neck cancers (HNSC) that show the highest levels of cell cycle activity in normal tissue, show the lowest baseline corrected cell cycle levels in tumours. This raises the intriguing question of whether there is a ceiling on cell cycle activity. Is it the case that driver gene mutations in HNSC are unable to push the cell cycle limits further as cells are already cycling at close to their maximum level? By extension, OV and UCS start from such a low level of activity that genomic aberrations (or exposure to sex hormones) have the scope to push cell cycle activity much higher than its starting point. If indeed cancer cells do have an upper limit on their cell cycle or proliferative capabilities, what is the limiting factor? One could speculate that cellular plasticity may hold the answer to these questions. Recent evidence has recognised cellular plasticity or phenotype switching as an essential process in disease (for a review of cell plasticity in cancer cells see here(24)), and one which takes many forms in cancer including epithelial to mesenchymal (EMT) transition(25), dedifferentiation(26), and transient spatial organization(27). While the experimental assessment of these processes lies beyond the scope of the work described herein, Malta *et al.* provide some evidence of the link between the cell cycle and cellular plasticity at a pan-cancer level. Using machine learning-derived stemness indices that were highly correlated to EMT markers, they showed that higher indices were associated with more proliferative breast cancer subgroups and with head and neck cancers relative to lower indices in kidney renal papillary cell carcinoma(28). It should be noted however that CESC, OV and UCS were of intermediate stemness, implying that stemness is not the only factor relevant to a theoretical upper ceiling of cell cycle activity.

Our analysis also showed a moderate to strong correlation (Spearman’s Rho = 0.61) between the BC-CCS and the *C19orf57* gene in Groups 1 and 2, and a moderate one (Spearman’s Rho = 0.32) in the rest of the pan-cancer cohort. Also called break repair meiotic recombinase recruitment factor 1 (*BRME1*), this gene has been shown by Zhang *et al.* to impair the mitotic *BRCA2*-*RAD51* homologous repair (HR) function in cancer cells and to be upregulated in brain and cervical cancers relative to paired normal tissues(29). Based on these findings, it could be speculated that through upregulation of *C19orf57* and subsequent sequestration of *BRCA2*, tumours promote genome instability and loss of strict control over the cell cycle. The strength of the correlations we saw in the Group 1 vs. 2 analysis relative to the one observed in rest of the cohort however suggests that this is unlikely to be the only factor driving large differences in baseline corrected cell cycle activity. Relatedly, we also saw a number of statistically significant differences in chromosomal arm-level amplifications and deletions in our Group 1 vs. 2 analyses that could be candidates also driving relative cell cycle activity. Their role at a pan-cancer level is less clear though as clustering on their basis showed no pattern of separation into low or high BC-CCS (Supplemental Figure 4).

The limitations of our study are as follows: First, the samples in this study are not matched tumour and normal tissue from the same patient, this means that we need a large sample size within each normal tissue type to derive the most accurate median values. Whilst this is true for many of the tissue types, cervical tissue (N = 13) stands out as one where caution should be taken with over-interpretation of the results. Second, we study broad chromosomal gains and losses rather than gene-centric copy number changes – this comes from our previous experience with this data and wanting to avoid a situation where the most changed genes in Group 2 would all come from the same chromosomal location. Third, we are focusing on cell cycle activity changes by applying the CCS signature to mRNA data extracted from an entire tumour sample as opposed to single cells. This means we get an average signal across all tumour cells and that the variance that would be seen in cell cycle activity at a single-cell level is not taken into account. The main strengths are: First, we apply a novel methodology to broadly reclassify tumour cell cycle activity in terms of its tissue of origin in order to derive basic biological insight. Second, we use multiple ‘omics data types to assess and further understand the differences between tumours of low and high BC-CCS; and third, we describe a new hypothesis of cell cycle activity having a ceiling that tumours may be unable to push past.

In summary, this study describes the reclassification of tumour cell cycle activity in terms of its normal tissue of origin. We show that in general, gynaecological cancers show the largest change in this activity and that it is likely driven by sex hormones, chromosomal arm-level alterations, and individual gene expression differences. Finally, we propose a new hypothesis of there being an upper-limit or “ceiling” on cell cycle activity in tumours at a pan-cancer level.

## Abbreviations

ACC: Adrenocortical carcinoma
AR: Androgen receptor
BC-CCS: Baseline Corrected-Cell Cycle Score
FDR: Benjamini-Hochberg correction
BLCA: Bladder Urothelial Carcinoma
LGG: Brain lower grade Glioma
BRME1: Break repair meiotic recombinase recruitment factor 1
BRCA: Breast invasive carcinoma
CCS: Cell Cycle Score
CESC: Cervical squamous cell carcinoma and endocervical adenocarcinoma
COAD: Colon adenocarcinoma
CDK: Cyclin dependent kinase
EMT: Epithelial to mesenchymal transition
ESCA: Esophageal carcinoma
ER-α: Estrogen receptor alpha
GSEA: Gene Set Enrichment Analyses
GDC: Genome Data Commons
GTEx: Genotype-Tissue Expression
GBM: Glioblastoma multiforme
HNSC: Head and Neck squamous cell carcinoma
HR: Homologous repair
HGNC: HUGO gene nomenclature committee
KICH: Kidney Chromophobe
KIRC: Kidney renal clear cell carcinoma
KIRP: Kidney renal papillary cell carcinoma
KEGG: Kyoto Encyclopedia of Genes and Genomes
LICH: Liver hepatocellular carcinoma
LUAD: Lung adenocarcinoma
LUSC: Lung squamous cell carcinoma
NIH: National Institute of Health
OV: Ovarian serous cystadenocarcinoma
PAAD: Pancreatic adenocarcinoma
PCA: Principal Component Analysis
PR: Progesterone receptor
PRAD: Prostate adenocarcinoma
SKCM: Skin Cutaneous Melanoma
STAD: Stomach adenocarcinoma
TGCT: Testicular Germ Cell tumours
TCGA: The Cancer Genome Atlas
TARGET: Therapeutically Applicable Research To Generate Effective Treatments
THCA: Thyroid carcinoma
Toil: The UCSC Toil RNAseq Recompute Compendium
UMAP: Uniform Manifold Approximation and Projection
UCS: Uterine Carcinosarcoma
UCEC: Uterine Corpus Endometrial Carcinoma

## Acknowledgments

None

## Funding

This work was supported by the Iris, Stig och Gerry Castenbäcks Stiftelse for cancer research (N.P.T.); the King Gustaf V Jubilee Foundation (N.P.T.); the Stockholm Cancer Society (Cancerföreningen i Stockholm to L.S.L.); the Swedish Cancer Society (Cancerfonden, N.P.T. grant number: 200802; L.S.L. grant number: 190140); the Swedish Research Council (Vetenskapsrådet, grant number 2020-02466 to L.S.L); the Swedish Research Council for Health, Working life and Welfare, (FORTE, grant number 2019-00477 to L.S.L.); ALF medicine (grant number LS2018-1157 to L.S.L.) and the Gösta Milton Donation Fund (Stiftelsen Gösta Miltons donationsfond, to L.S.L.

## Authors’ contributions

AL, JJJY, LSL and NPT contributed the study concept and design. AL, JJJY and NPT contributed to the acquisition and analyses of data. All authors interpreted the data, drafted the manuscript, read and approved the final version.

## Ethics approval and consent to participate

Publicly available data - Not applicable

## Consent for publication

Publicly available data - Not applicable

## Data deposition and code

The data used in this study are publicly available on National Institute of Health (NIH) (https://gdc.cancer.gov/about-data/publications/pancanatlas) and UCSC Xena website (http://xena.ucsc.edu). R-code to reproduce the results of this study are publicly available at https://github.com/arianlundberg/BC-CCS.

## Competing Interests

None.

## Notes

### Competing Interest Statement

The authors have declared no competing interest.

