## Supplemental Figure 1. for "Reclassifying Cancer: Defining tumour cell cycle activity in terms of its tissue of origin in over 13,000 samples"

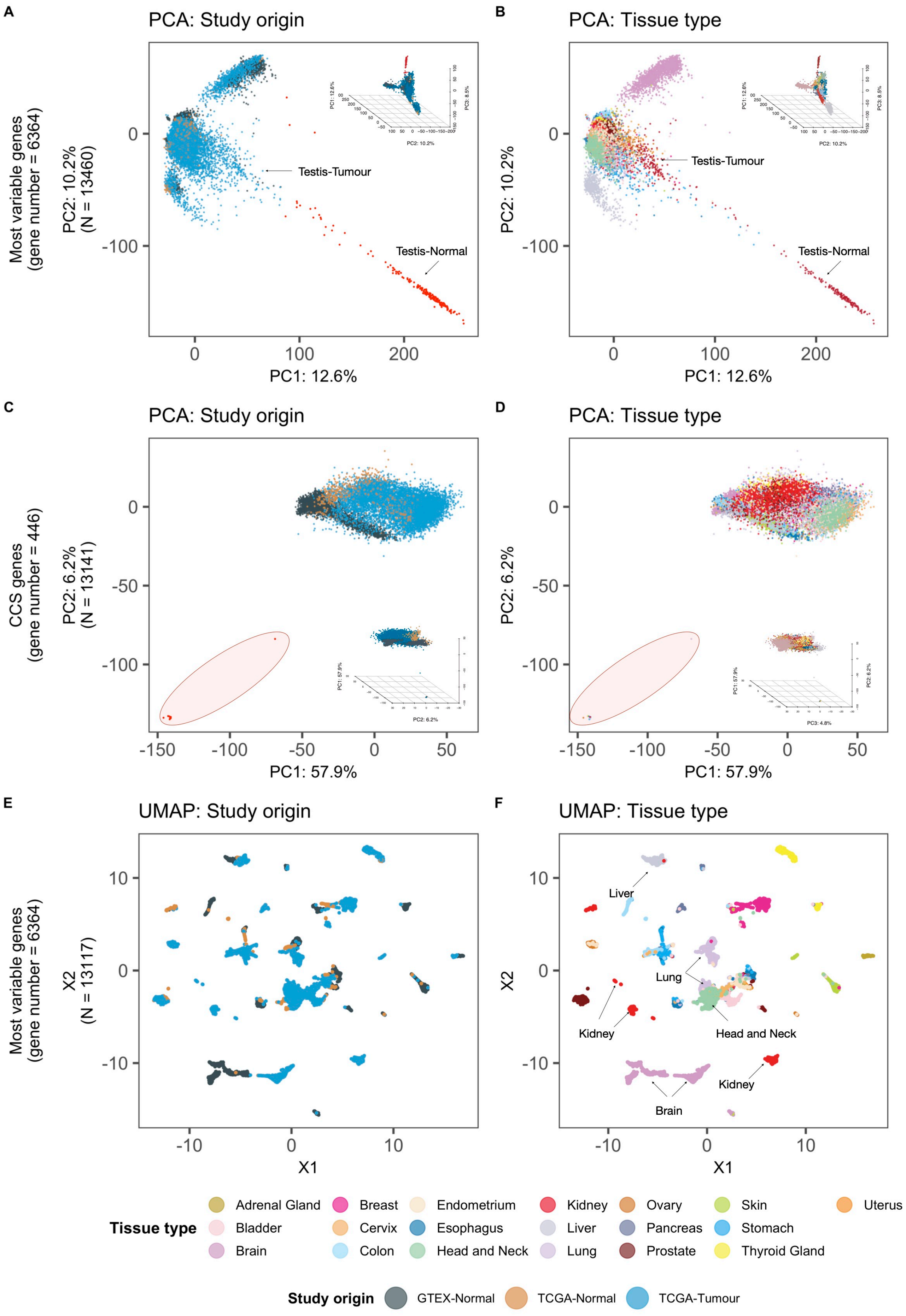


**Supplemental Figure 1. Dimensionality reduction of Pan-Cancer and GTEx data**

Principle component analyses (PCA) using most variable genes in the data: A) based on study origin and B) tissue types; PCA plots of the data using genes incorporated in the Cell cycle score (CCS): C) based on study origin and D) tissue types. Uniform Manifold Approximation and Projection (UMAP) plots of the data representing the clusters of tumors: based on E) Tissue type and F) Study origin after exclusion of the outliers.
