## Supplemental Figure 2. for "Reclassifying Cancer: Defining tumour cell cycle activity in terms of its tissue of origin in over 13,000 samples"

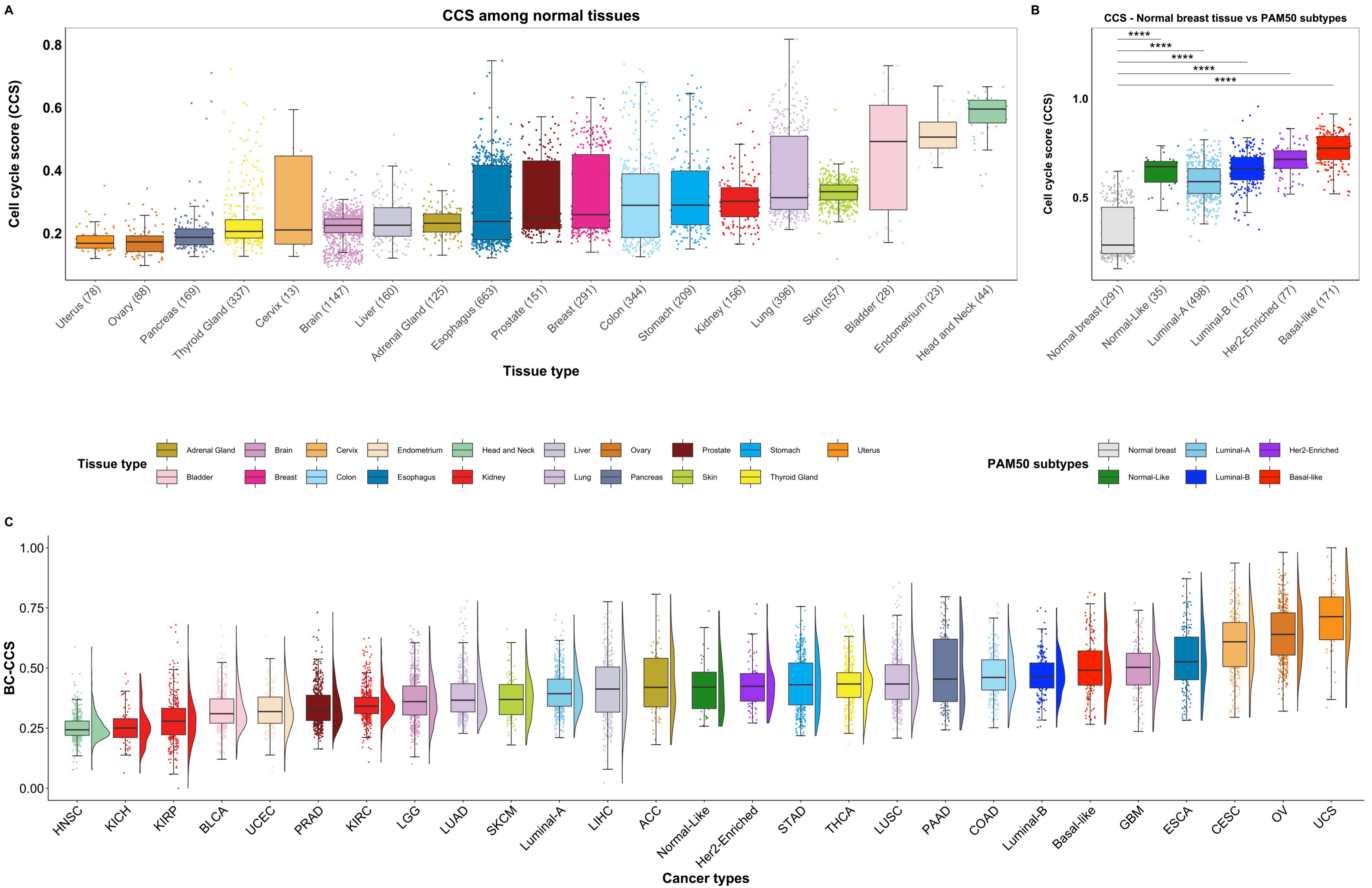


**Supplemental Figure 2. Cell cycle activity among normal tissues and PAN-Cancer samples.**
A) Boxplots representing Cell cycle score (CCS) as a surrogate for cell cycle activity among normal tissues. B) CCS score range among breast cancer PAM50 subtypes C) Boxplots and violin plots showing the Baseline Corrected - Cell Cycle Score (BC-CCS) among different tumour types including PAM50 subtypes of breast cancer tumours.
