## Supplemental Figure 3. for "Reclassifying Cancer: Defining tumour cell cycle activity in terms of its tissue of origin in over 13,000 samples"

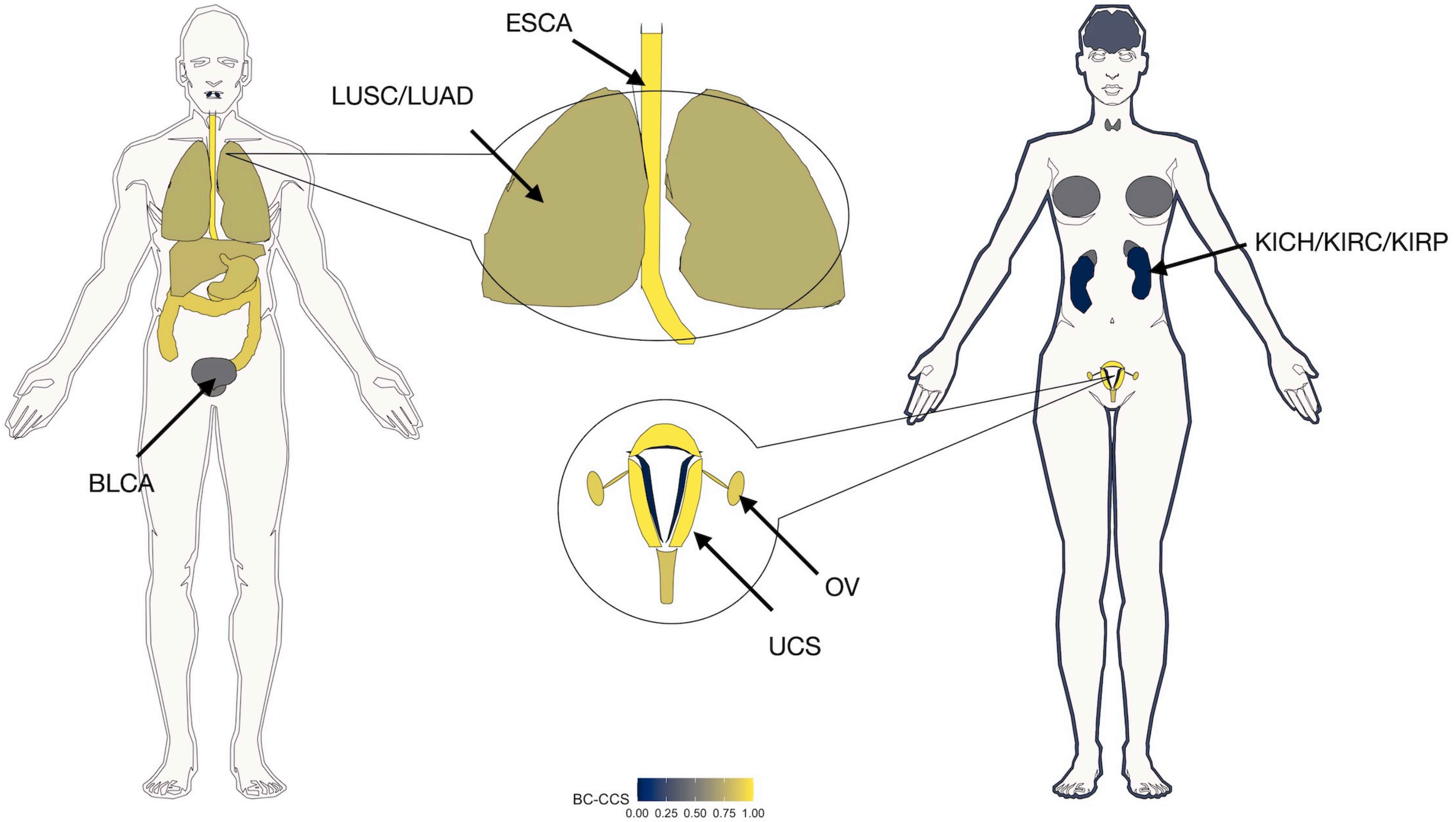


**Supplemental Figure 3. Baseline Corrected - Cell Cycle Score (BC-CCS) at PAN-Cancer level**

An anatomical visualization of BC-CCS among PAN-Cancer. Bladder Urothelial Carcinoma (BLCA), Esophageal carcinoma (ESCA), Kidney Chromophobe (KICH), Kidney renal clear cell carcinoma (KIRC), Kidney renal papillary cell carcinoma (KIRP), Lung adenocarcinoma (LUAD), Lung squamous cell carcinoma (LUSC), Ovarian serous cystuadenocarcinoma (OV) and Uterine Carcinosarcoma (UCS).
