## Supplemental Figure 4. for "Reclassifying Cancer: Defining tumour cell cycle activity in terms of its tissue of origin in over 13,000 samples"

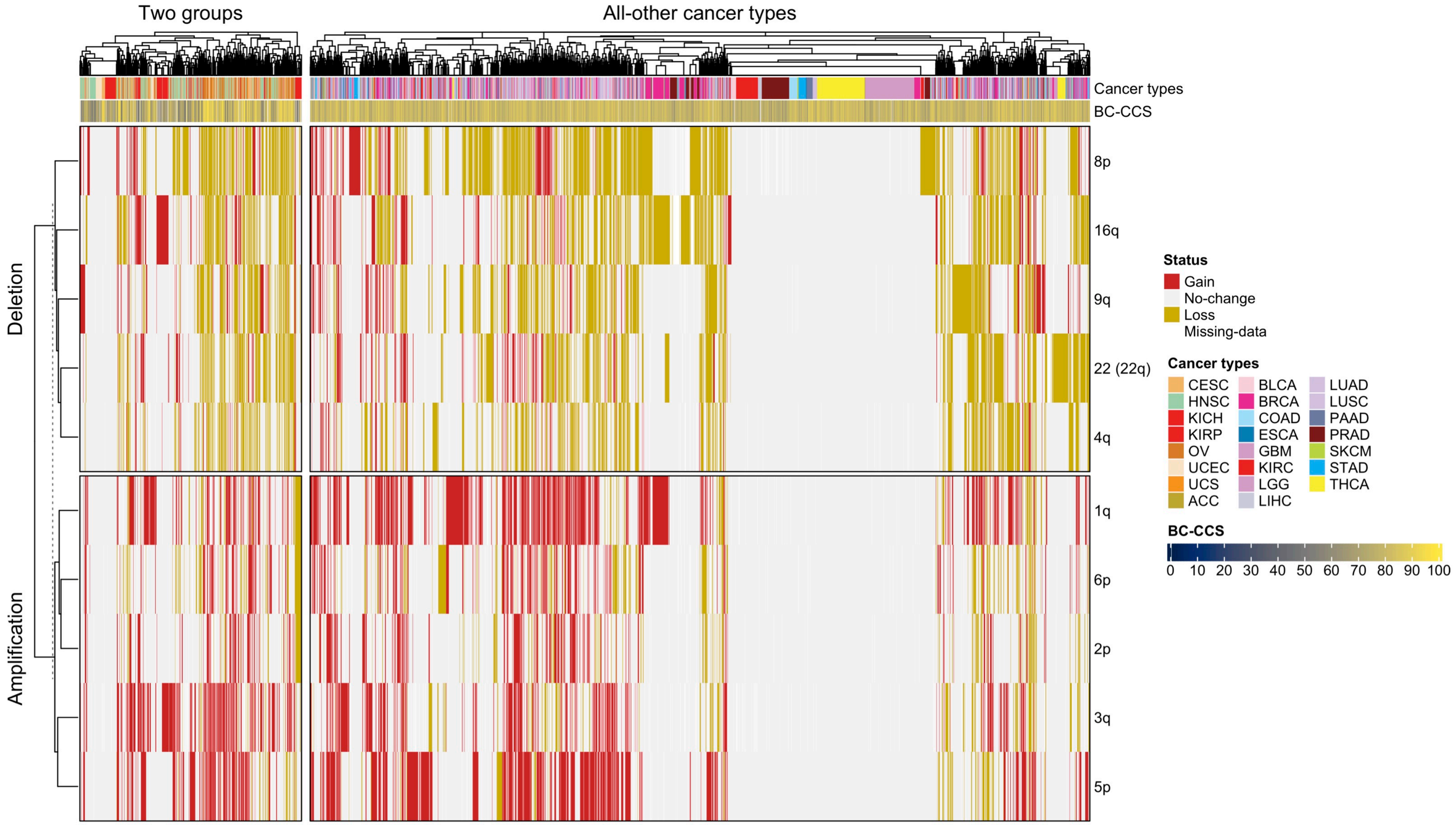


**Supplemental Figure 4. Heatmap of aneuploidy status at PAN-Cancer level.**
A heatmap representing the amplification and deletion status of tumors at top 5 chromosomal locations with highest Gain/Loss in Group 2 relative to Group 1 as shown in Figure 4D and E, respectively. BC-CCS: Baseline Corrected - Cell Cycle Score.
