## Supplemental Figure 5. for "Reclassifying Cancer: Defining tumour cell cycle activity in terms of its tissue of origin in over 13,000 samples"

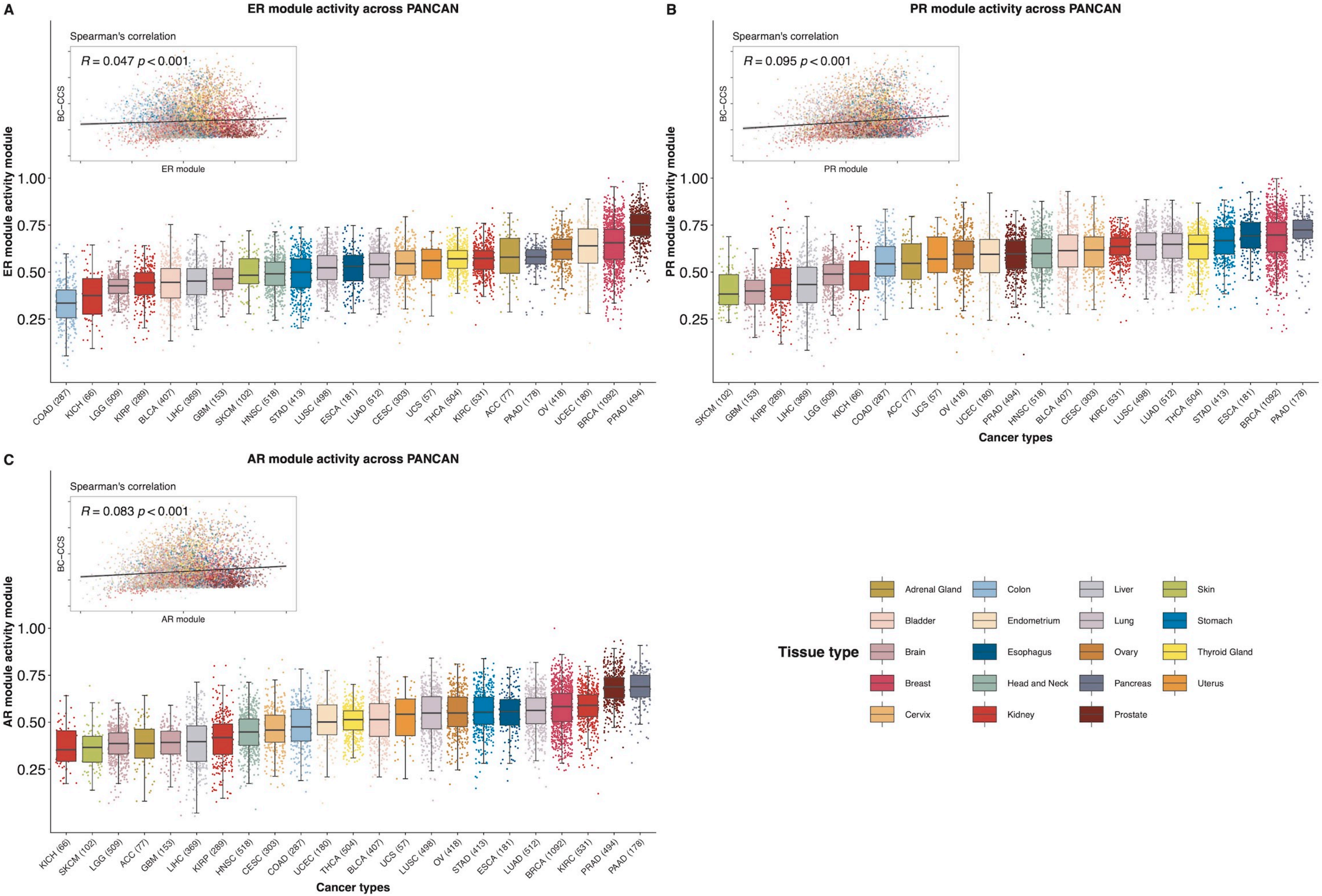


**Supplemental Figure 5. Estrogen, Progesterone and Androgen gene modules activity across PANCAN**

Box plots representing the A) Estrogen (ER), B) Progesterone (PR), C) Androgen (AR) gene modules activity across PANCAN. Spearman's correlation *R* shows the correlation between Baseline Corrected - Cell Cycle Score (BC-CCS) and ER,PR and AR modules.
