## Supplemental Table 1. for "Reclassifying Cancer: Defining tumour cell cycle activity in terms of its tissue of origin in over 13,000 samples"

Supplemental table 1.
Cell Cycle score genes (CCS)

| EntrezID | HGNC Symbol | Chromosomal location | Approved Name | Included in the study |
|---|---|---|---|---|
| 10057 | ABCC5 | 3q27 | ATP-binding cassette, sub-family C (CFTR/MRP), member 5 | Yes |
| 25 | ABL1 | 9q34.1 | ABL proto-oncogene 1, non-receptor tyrosine kinase | Yes |
| 10097 | ACTR2 | 2p14 | ARP2 actin-related protein 2 homolog (yeast) | Yes |
| 127 | ADH4 | 4q22 | alcohol dehydrogenase 4 (class II), pi polypeptide | Yes |
| 60312 | AFAP1 | 4p16 | actin filament associated protein 1 | Yes |
| 54806 | AHI1 | 6q23.2 | Abelson helper integration site 1 | Yes |
| 55122 | AKIRIN2 | 6q15 | akirin 2 | Yes |
| 84266 | ALKBH7 | 19p13.3 | alkB, alkylation repair homolog 7 (E. coli) | Yes |
| 64682 | ANAPC1 | 2q12.1 | anaphase promoting complex subunit 1 | Yes |
| 10393 | ANAPC10 | 4q31 | anaphase promoting complex subunit 10 | Yes |
| 51529 | ANAPC11 | 17q25.3 | anaphase promoting complex subunit 11 | Yes |
| 25847 | ANAPC13 | 3q22.1 | anaphase promoting complex subunit 13 | Yes |
| 29882 | ANAPC2 | 9q34.3 | anaphase promoting complex subunit 2 | Yes |
| 29945 | ANAPC4 | 4p15.31 | anaphase promoting complex subunit 4 | Yes |
| 51433 | ANAPC5 | 12q24.31 | anaphase promoting complex subunit 5 | Yes |
| 51434 | ANAPC7 | 12q13.12 | anaphase promoting complex subunit 7 | Yes |
| 55608 | ANKRD10 | 13q33.3 | ankyrin repeat domain 10 | Yes |
| 400986 | ANKRD36C | 2q11.1 | ankyrin repeat domain 36C | Yes |
| 54443 | ANLN | 7p15-p14 | anillin, actin binding protein | Yes |
| 84168 | ANTXR1 | 2p13.1 | anthrax toxin receptor 1 | Yes |
| 314 | AOC2 | 17q21 | amine oxidase, copper containing 2 (retina-specific) | Yes |
| 375 | ARF1 | 1q42.13 | ADP-ribosylation factor 1 | Yes |
| 55082 | ARGLU1 | 13q33.3 | arginine and glutamate rich 1 | Yes |
| 9824 | ARHGAP11A | 15q13.3 | Rho GTPase activating protein 11A | Yes |
| 89839 | ARHGAP11B | 15q13.2 | Rho GTPase activating protein 11B | No |
| 84904 | ARHGEF39 | 9p13.3 | Rho guanine nucleotide exchange factor (GEF) 39 | Yes |
| 10124 | ARL4A | 7p21.3 | ADP-ribosylation factor-like 4A | Yes |
| 23204 | ARL6IP1 | 16p12-p11.2 | ADP-ribosylation factor-like 6 interacting protein 1 | Yes |
| 55723 | ASF1B | 19p13.12 | anti-silencing function 1B histone chaperone | Yes |
| 434 | ASIP | 20q11.2-q12 | agouti signaling protein | Yes |
| 171023 | ASXL1 | 20q11 | additional sex combs like transcriptional regulator 1 | Yes |
| 29028 | ATAD2 | 8q24.13 | ATPase family, AAA domain containing 2 | Yes |
| 64225 | ATL2 | 2p22.3 | atlastin GTPase 2 | Yes |
| 472 | ATM | 11q22-q23 | ATM serine/threonine kinase | Yes |
| 545 | ATR | 3q23 | ATR serine/threonine kinase | Yes |
| 84126 | ATRIP | 3p21.31 | ATR interacting protein | Yes |
| 6790 | AURKA | 20q13 | aurora kinase A | Yes |
| 9212 | AURKB | 17p13.1 | aurora kinase B | Yes |
| 329 | BIRC2 | 11q22 | baculoviral IAP repeat containing 2 | Yes |
| 330 | BIRC3 | 11q22 | baculoviral IAP repeat containing 3 | Yes |
| 332 | BIRC5 | 17q25.3 | baculoviral IAP repeat containing 5 | Yes |
| 650 | BMP2 | 20p12 | bone morphogenetic protein 2 | Yes |
| 79866 | BORA | 13q22.1 | bora, aurora kinase A activator | Yes |
| 672 | BRCA1 | 17q21.31 | breast cancer 1, early onset | Yes |
| 10902 | BRD8 | 5q31 | bromodomain containing 8 | Yes |
| 22903 | BTBD3 | 20p12.2 | BTB (POZ) domain containing 3 | Yes |
| 153579 | BTNL9 | 5q35.3 | butyrophilin-like 9 | Yes |
| 699 | BUB1 | 2q13 | BUB1 mitotic checkpoint serine/threonine kinase | Yes |
| 701 | BUB1B | 15q15 | BUB1 mitotic checkpoint serine/threonine kinase B | Yes |
| 9184 | BUB3 | 10q24 | BUB3 mitotic checkpoint protein | Yes |
| 65250 | C5orf42 | 5p13.2 | chromosome 5 open reading frame 42 | No |
| 836 | CASP3 | 4q34 | caspase 3, apoptosis-related cysteine peptidase | Yes |
| 203260 | CCDC107 | 9q13.3 | coiled-coil domain containing 107 | Yes |
| 8900 | CCNA1 | 13q12.3-q13 | cyclin A1 | Yes |
| 890 | CCNA2 | 4q27 | cyclin A2 | Yes |
| 891 | CCNB1 | 5q12 | cyclin B1 | Yes |
| 9133 | CCNB2 | 15q21.3 | cyclin B2 | Yes |
| 85417 | CCNB3 | Xp11 | cyclin B3 | Yes |
| 595 | CCND1 | 11q13 | cyclin D1 | Yes |
| 894 | CCND2 | 12p13 | cyclin D2 | Yes |
| 896 | CCND3 | 6p21 | cyclin D3 | Yes |
| 898 | CCNE1 | 19q12 | cyclin E1 | Yes |
| 9134 | CCNE2 | 8q22.1 | cyclin E2 | Yes |
| 899 | CCNF | 16p13.3 | cyclin F | Yes |
| 902 | CCNH | 5q13.3-q14 | cyclin H | Yes |
| 8556 | CDC14A | 1p21 | cell division cycle 14A | Yes |
| 8555 | CDC14B | 9q22.3 | cell division cycle 14B | Yes |
| 8881 | CDC16 | 13q34 | cell division cycle 16 | Yes |

| | | | | |
|---|---|---|---|---|
| 991 | CDC20 | 1p34.1 | cell division cycle 20 | Yes |
| 8697 | CDC23 | 5q31 | cell division cycle 23 | Yes |
| 993 | CDC25A | 3p21 | cell division cycle 25A | Yes |
| 994 | CDC25B | 20p13 | cell division cycle 25B | Yes |
| 995 | CDC25C | 5q31 | cell division cycle 25C | Yes |
| 246184 | CDC26 | 9q32 | cell division cycle 26 | Yes |
| 996 | CDC27 | 17q21.32 | cell division cycle 27 | Yes |
| 8318 | CDC45 | 22q11.21 | cell division cycle 45 | Yes |
| 990 | CDC6 | 17q21.3 | cell division cycle 6 | Yes |
| 8317 | CDC7 | 1p22 | cell division cycle 7 | Yes |
| 157313 | CDCA2 | 8p21.2 | cell division cycle associated 2 | Yes |
| 83461 | CDCA3 | 12p13.31 | cell division cycle associated 3 | Yes |
| 113130 | CDCA5 | 11q13.1 | cell division cycle associated 5 | Yes |
| 83879 | CDCA7 | 2q31.1 | cell division cycle associated 7 | Yes |
| 55536 | CDCA7L | 7p15.3 | cell division cycle associated 7-like | Yes |
| 55143 | CDCA8 | 1p34.3 | cell division cycle associated 8 | Yes |
| 999 | CDH1 | 16q22.1 | cadherin 1, type 1, E-cadherin (epithelial) | Yes |
| 64403 | CDH24 | 14q11.2 | cadherin 24, type 2 | Yes |
| 983 | CDK1 | 10q21.2 | cyclin-dependent kinase 1 | Yes |
| 1017 | CDK2 | 12q13 | cyclin-dependent kinase 2 | Yes |
| 1019 | CDK4 | 12q13 | cyclin-dependent kinase 4 | Yes |
| 1021 | CDK6 | 7q21-q22 | cyclin-dependent kinase 6 | Yes |
| 1022 | CDK7 | 5q12.1 | cyclin-dependent kinase 7 | Yes |
| 6792 | CDKL5 | Xp22 | cyclin-dependent kinase-like 5 | Yes |
| 1026 | CDKN1A | 6p21.1 | cyclin-dependent kinase inhibitor 1A (p21, Cip1) | Yes |
| 1027 | CDKN1B | 12p13.1-p12 | cyclin-dependent kinase inhibitor 1B (p27, Kip1) | Yes |
| 1028 | CDKN1C | 11p15.5 | cyclin-dependent kinase inhibitor 1C (p57, Kip2) | Yes |
| 1029 | CDKN2A | 9p21 | cyclin-dependent kinase inhibitor 2A | Yes |
| 55602 | CDKN2AIP | 4q35.1 | CDKN2A interacting protein | Yes |
| 1030 | CDKN2B | 9p21 | cyclin-dependent kinase inhibitor 2B (p15, inhibits CDK4) | Yes |
| 1031 | CDKN2C | 1p32.3 | cyclin-dependent kinase inhibitor 2C (p18, inhibits CDK4) | Yes |
| 1032 | CDKN2D | 19p13 | cyclin-dependent kinase inhibitor 2D (p19, inhibits CDK4) | Yes |
| 1033 | CDKN3 | 14q22 | cyclin-dependent kinase inhibitor 3 | Yes |
| 1058 | CENPA | 2p23.3 | centromere protein A | Yes |
| 1062 | CENPE | 4q24-q25 | centromere protein E, 312kDa | Yes |
| 1063 | CENPF | 1q41 | centromere protein F, 350/400kDa | Yes |
| 91687 | CENPL | 1q25.1 | centromere protein L | Yes |
| 55166 | CENPQ | 6p12.3 | centromere protein Q | Yes |
| 79682 | CENPU | 4q35.1 | centromere protein U | Yes |
| 55165 | CEP55 | 10q24.1 | centrosomal protein 55kDa | Yes |
| 80321 | CEP70 | 3q22.3 | centrosomal protein 70kDa | Yes |
| 8208 | CHAF1B | 21q22.2 | chromatin assembly factor 1, subunit B (p60) | Yes |
| 1111 | CHEK1 | 11q24.2 | checkpoint kinase 1 | Yes |
| 11200 | CHEK2 | 22q12.1 | checkpoint kinase 2 | Yes |
| 11113 | CIT | 12q24.23 | citron rho-interacting serine/threonine kinase | Yes |
| 26586 | CKAP2 | 13q14 | cytoskeleton associated protein 2 | Yes |
| 150468 | CKAP2L | 2q13 | cytoskeleton associated protein 2-like | Yes |
| 9793 | CKAP5 | 11p11.2 | cytoskeleton associated protein 5 | Yes |
| 1163 | CKS1B | 1q21.2 | CDC28 protein kinase regulatory subunit 1B | Yes |
| 1164 | CKS2 | 9q22 | CDC28 protein kinase regulatory subunit 2 | Yes |
| 63967 | CLSPN | 1p34.3 | claspin | Yes |
| 29097 | CNIH4 | 1q42.12 | cornichon family AMPA receptor auxiliary protein 4 | Yes |
| 116840 | CNTROB | 17p13.1 | centrobin, centrosomal BRCA2 interacting protein | Yes |
| 51004 | COQ6 | 14q24.1 | coenzyme Q6 monooxygenase | Yes |
| 1387 | CREBBP | 16p13.3 | CREB binding protein | Yes |
| 58487 | CREBZF | 11q14.1 | CREB/ATF bZIP transcription factor | Yes |
| 1495 | CTNNA1 | 5q31.2 | catenin (cadherin-associated protein), alpha 1, 102kDa | Yes |
| 8454 | CUL1 | 7q36.1 | cullin 1 | Yes |
| 285440 | CYP4V2 | 4q35.2 | cytochrome P450, family 4, subfamily V, polypeptide 2 | Yes |
| 10926 | DBF4 | 7q21.3 | DBF4 zinc finger | Yes |
| 79077 | DCTPP1 | 16p11.2 | dCTP pyrophosphatase 1 | Yes |
| 23586 | DDX58 | 9p12 | DEAD (Asp-Glu-Ala-Asp) box polypeptide 58 | Yes |
| 55635 | DEPDC1 | 1p31.2 | DEP domain containing 1 | Yes |
| 55789 | DEPDC1B | 5q12 | DEP domain containing 1B | Yes |
| 200895 | DHFRL1 | 3q11.2 | dihydrofolate reductase-like 1 | No |
| 1736 | DKC1 | Xq28 | dyskeratosis congenita 1, dyskerin | Yes |
| 9787 | DLGAP5 | 14q22.3 | discs, large (Drosophila) homolog-associated protein 5 | Yes |
| 8701 | DNAH11 | 7p21 | dynein, axonemal, heavy chain 11 | Yes |
| 3337 | DNAJB1 | 19p13.12 | DnaJ (Hsp40) homolog, subfamily B, member 1 | Yes |
| 10049 | DNAJB6 | 7q36.3 | DnaJ (Hsp40) homolog, subfamily B, member 6 | Yes |
| 1789 | DNMT3B | 20q11.2 | DNA (cytosine-5-)-methyltransferase 3 beta | Yes |
| 29980 | DONSON | 21q22.1 | downstream neighbor of SON | Yes |

| | | | | |
|---|---|---|---|---|
| 79075 | DSCC1 | 8q24.12 | DNA replication and sister chromatid cohesion 1 | Yes |
| 51514 | DTL | 1q32 | denticleless E3 ubiquitin protein ligase homolog (Drosophila) | Yes |
| 8655 | DYNLL1 | 12q24.23 | dynein, light chain, LC8-type 1 | Yes |
| 1869 | E2F1 | 20q11 | E2F transcription factor 1 | Yes |
| 1870 | E2F2 | 1p36 | E2F transcription factor 2 | Yes |
| 1871 | E2F3 | 6p22 | E2F transcription factor 3 | Yes |
| 1874 | E2F4 | 16q22.1 | E2F transcription factor 4, p107/p130-binding | Yes |
| 1875 | E2F5 | 8q21.2 | E2F transcription factor 5, p130-binding | Yes |
| 1876 | E2F6 | 2p25.1 | E2F transcription factor 6 | Yes |
| 79733 | E2F8 | 11p15 | E2F transcription factor 8 | Yes |
| 1894 | ECT2 | 3q26.1-q26.2 | epithelial cell transforming 2 | Yes |
| 114327 | EFHC1 | 6p12.3 | EF-hand domain (C-terminal) containing 1 | Yes |
| 56648 | EIF5A2 | 3q26.2 | eukaryotic translation initiation factor 5A2 | Yes |
| 2033 | EP300 | 22q13.2 | E1A binding protein p300 | Yes |
| 10595 | ERN2 | 16p12.2 | endoplasmic reticulum to nucleus signaling 2 | Yes |
| 157570 | ESCO2 | 8p21.1 | establishment of sister chromatid cohesion N-acetyltransferase 2 | Yes |
| 9700 | ESPL1 | 12q13.13 | extra spindle pole bodies homolog 1 (S. cerevisiae) | Yes |
| 2118 | ETV4 | 17q21 | ets variant 4 | Yes |
| 9156 | EXO1 | 1q43 | exonuclease 1 | Yes |
| 2146 | EZH2 | 7q35-q36 | enhancer of zeste 2 polycomb repressive complex 2 subunit | Yes |
| 284611 | FAM102B | 1p13.3 | family with sequence similarity 102, member B | Yes |
| 374393 | FAM111B | 11q12.1 | family with sequence similarity 111, member B | Yes |
| 10712 | FAM189B | 1q21 | family with sequence similarity 189, member B | Yes |
| 56204 | FAM214A | 15q21.2-q21.3 | family with sequence similarity 214, member A | Yes |
| 29902 | FAM216A | 12q24.11 | family with sequence similarity 216, member A | Yes |
| 54478 | FAM64A | 17p13.2 | family with sequence similarity 64, member A | No |
| 729533 | FAM72A | 1q32.1 | family with sequence similarity 72, member A | Yes |
| 653820 | FAM72B | 1p12 | family with sequence similarity 72, member B | Yes |
| 81610 | FAM83D | 20q11.23 | family with sequence similarity 83, member D | Yes |
| 157638 | FAM84B | 8q24.13 | family with sequence similarity 84, member B | No |
| 2177 | FANCD2 | 3p25.3 | Fanconi anemia, complementation group D2 | Yes |
| 55215 | FANCI | 15q26.1 | Fanconi anemia, complementation group I | Yes |
| 2237 | FEN1 | 11q12 | flap structure-specific endonuclease 1 | Yes |
| 123811 | FOPNL | 16p13.11 | FGFR1OP N-terminal like | Yes |
| 2305 | FOXM1 | 12p13 | forkhead box M1 | Yes |
| 51343 | FZR1 | 19p13.3 | fizzy/cell division cycle 20 related 1 (Drosophila) | Yes |
| 55632 | G2E3 | 14q12 | G2/M-phase specific E3 ubiquitin protein ligase | Yes |
| 2553 | GABPB1 | 15q21.2 | GA binding protein transcription factor, beta subunit 1 | Yes |
| 1647 | GADD45A | 1p31.2 | growth arrest and DNA-damage-inducible, alpha | Yes |
| 4616 | GADD45B | 19p13.3 | growth arrest and DNA-damage-inducible, beta | Yes |
| 10912 | GADD45G | 9q22.1-q22.2 | growth arrest and DNA-damage-inducible, gamma | Yes |
| 2619 | GAS1 | 9q21.3-q22 | growth arrest-specific 1 | Yes |
| 283431 | GAS2L3 | 12q23.1 | growth arrest-specific 2 like 3 | Yes |
| 2621 | GAS6 | 13q34 | growth arrest-specific 6 | Yes |
| 2730 | GCLM | 1p21 | glutamate-cysteine ligase, modifier subunit | Yes |
| 257144 | GCSAM | 3q13.13 | germinal center-associated, signaling and motility | Yes |
| 51659 | GINS2 | 16q24.1 | GINS complex subunit 2 (Psf2 homolog) | Yes |
| 51053 | GMNN | 6p21.32 | geminin, DNA replication inhibitor | Yes |
| 23015 | GOLGA8A | 15q14 | golgin A8 family, member A | Yes |
| 29899 | GPSM2 | 1p13.3 | G-protein signaling modulator 2 | Yes |
| 80273 | GRPEL1 | 4p16 | GrpE-like 1, mitochondrial (E. coli) | Yes |
| 23199 | GSE1 | 16q24.1 | Gse1 coiled-coil protein | Yes |
| 2932 | GSK3B | 3q13.3 | glycogen synthase kinase 3 beta | Yes |
| 51512 | GTSE1 | 22q13.2-q13.3 | G-2 and S-phase expressed 1 | Yes |
| 3005 | H1F0 | 22q13.1 | H1 histone family, member 0 | No |
| 3014 | H2AFX | 11q23.3 | H2A histone family, member X | No |
| 3048 | HBG2 | 11p15.5 | hemoglobin, gamma G | Yes |
| 3065 | HDAC1 | 1p34 | histone deacetylase 1 | Yes |
| 3066 | HDAC2 | 6q21 | histone deacetylase 2 | Yes |
| 8841 | HDAC3 | 5q31.1-q31.2 | histone deacetylase 3 | Yes |
| 9759 | HDAC4 | 2q37.3 | histone deacetylase 4 | Yes |
| 10014 | HDAC5 | 17q21 | histone deacetylase 5 | Yes |
| 10013 | HDAC6 | Xp11.23 | histone deacetylase 6 | Yes |
| 55869 | HDAC8 | Xq13 | histone deacetylase 8 | Yes |
| 3070 | HELLS | 10q24.2 | helicase, lymphoid-specific | Yes |
| 3091 | HIF1A | 14q23.2 | inducible factor 1, alpha subunit (basic helix-loop-helix transcriptio | Yes |
| 135114 | HINT3 | 6q22.33 | histidine triad nucleotide binding protein 3 | Yes |
| 8334 | HIST1H2AC | 6p22.1 | histone cluster 1, H2ac | No |
| 8364 | HIST1H4C | 6p22.1 | histone cluster 1, H4c | No |
| 8349 | HIST2H2BE | 1q21.2 | histone cluster 2, H2be | No |
| 55355 | HJURP | 2q37.1 | Holliday junction recognition protein | Yes |
| 3122 | HLA-DRA | 6p21.3 | major histocompatibility complex, class II, DR alpha | Yes |

| | | | | |
|---|---|---|---|---|
| 10362 | HMG20B | 19p13.3 | high mobility group 20B | Yes |
| 3148 | HMGB2 | 4q31 | high mobility group box 2 | Yes |
| 3149 | HMGB3 | Xq28 | high mobility group box 3 | Yes |
| 3161 | HMMR | 5q34 | hyaluronan-mediated motility receptor (RHAMM) | Yes |
| 51155 | HN1 | 17q25.1 | hematological and neurological expressed 1 | No |
| 84072 | HORMAD1 | 1q21.2 | HORMA domain containing 1 | Yes |
| 50809 | HP1BP3 | 1p36.12 | heterochromatin protein 1, binding protein 3 | Yes |
| 10247 | HRSP12 | 8q22 | heat-responsive protein 12 | No |
| 3305 | HSPA1L | 6p21.3 | heat shock 70kDa protein 1-like | Yes |
| 3306 | HSPA2 | 14q23 | heat shock 70kDa protein 2 | Yes |
| 3308 | HSPA4 | 5q31.1 | heat shock 70kDa protein 4 | Yes |
| 3312 | HSPA8 | 11q24.1 | heat shock 70kDa protein 8 | Yes |
| 26353 | HSPB8 | 12q24.23 | heat shock 22kDa protein 8 | Yes |
| 3620 | IDO1 | 8p12-p11 | indoleamine 2,3-dioxygenase 1 | Yes |
| 51278 | IER5 | 1q25.3 | immediate early response 5 | Yes |
| 3433 | IFIT2 | 10q23.31 | interferon-induced protein with tetratricopeptide repeats 2 | Yes |
| 3569 | IL6 | 7p21-p15 | interleukin 6 | Yes |
| 3575 | IL7R | 5p13 | interleukin 7 receptor | Yes |
| 51763 | INPP5K | 17p13.3 | inositol polyphosphate-5-phosphatase K | Yes |
| 51141 | INSIG2 | 2q14.1 | insulin induced gene 2 | Yes |
| 55656 | INTS8 | 8q22.1 | integrator complex subunit 8 | Yes |
| 128239 | IQGAP3 | 1q21.3 | IQ motif containing GTPase activating protein 3 | Yes |
| 3654 | IRAK1 | Xq28 | interleukin-1 receptor-associated kinase 1 | Yes |
| 10625 | IVNS1ABP | 1q25.1-q31.1 | influenza virus NS1A binding protein | Yes |
| 25948 | KBTBD2 | 7p14.3 | kelch repeat and BTB (POZ) domain containing 2 | Yes |
| 57650 | KIAA1524 | 3q13.13 | KIAA1524 | No |
| 3832 | KIF11 | 10q24.1 | kinesin family member 11 | Yes |
| 9928 | KIF14 | 1q32.1 | kinesin family member 14 | Yes |
| 9585 | KIF20B | 10q23.31 | kinesin family member 20B | Yes |
| 9493 | KIF23 | 15q23 | kinesin family member 23 | Yes |
| 11004 | KIF2C | 1p34.1 | kinesin family member 2C | Yes |
| 3799 | KIF5B | 10p11.22 | kinesin family member 5B | Yes |
| 3833 | KIFC1 | 6p21.32 | kinesin family member C1 | Yes |
| 1316 | KLF6 | 10p15 | Kruppel-like factor 6 | Yes |
| 687 | KLF9 | 9q21.11 | Kruppel-like factor 9 | Yes |
| 8564 | KMO | 1q42-q44 | kynurenine 3-monooxygenase (kynurenine 3-hydroxylase) | Yes |
| 90417 | KNSTRN | 15q15.1 | kinetochore-localized astrin/SPAG5 binding protein | Yes |
| 3838 | KPNA2 | 17q24.2 | karyopherin alpha 2 (RAG cohort 1, importin alpha 1) | Yes |
| 23367 | LARP1 | 5q33.2 | La ribonucleoprotein domain family, member 1 | Yes |
| 3930 | LBR | 1q42.1 | lamin B receptor | Yes |
| 200879 | LIPH | 3q27 | lipase, member H | Yes |
| 4000 | LMNA | 1q22 | lamin A/C | Yes |
| 55791 | LRIF1 | 1p13.3 | ligand dependent nuclear receptor interacting factor 1 | Yes |
| 10234 | LRRC17 | 7q22.1 | leucine rich repeat containing 17 | Yes |
| 4054 | LTBP3 | 11q12 | latent transforming growth factor beta binding protein 3 | Yes |
| 8379 | MAD1L1 | 7p22 | MAD1 mitotic arrest deficient-like 1 (yeast) | Yes |
| 4085 | MAD2L1 | 4q27 | MAD2 mitotic arrest deficient-like 1 (yeast) | Yes |
| 10459 | MAD2L2 | 1p36 | MAD2 mitotic arrest deficient-like 2 (yeast) | Yes |
| 5608 | MAP2K6 | 17q | mitogen-activated protein kinase kinase 6 | Yes |
| 5603 | MAPK13 | 6p21 | mitogen-activated protein kinase 13 | Yes |
| 84930 | MASTL | 10p12.1 | microtubule associated serine/threonine kinase-like | Yes |
| 154141 | MBOAT1 | 6p22.3 | membrane bound O-acyltransferase domain containing 1 | Yes |
| 55388 | MCM10 | 10p13 | minichromosome maintenance complex component 10 | Yes |
| 4171 | MCM2 | 3q21 | minichromosome maintenance complex component 2 | Yes |
| 4172 | MCM3 | 6p12 | minichromosome maintenance complex component 3 | Yes |
| 4173 | MCM4 | 8q12-q13 | minichromosome maintenance complex component 4 | Yes |
| 4174 | MCM5 | 22q13.1-q13.2 | minichromosome maintenance complex component 5 | Yes |
| 4175 | MCM6 | 2q14-q21 | minichromosome maintenance complex component 6 | Yes |
| 4176 | MCM7 | 7q21.3-q22.1 | minichromosome maintenance complex component 7 | Yes |
| 84515 | MCM8 | 20p12.3 | minichromosome maintenance complex component 8 | Yes |
| 254394 | MCM9 | 6q22.31 | minichromosome maintenance complex component 9 | Yes |
| 4193 | MDM2 | 12q13-q14 | MDM2 proto-oncogene, E3 ubiquitin protein ligase | Yes |
| 9833 | MELK | 9p13.1 | maternal embryonic leucine zipper kinase | Yes |
| 4221 | MEN1 | 11q13 | multiple endocrine neoplasia I | Yes |
| 56257 | MEPCE | 7q22.1 | methylphosphate capping enzyme | Yes |
| 4247 | MGAT2 | 14q21 | ıosyl (alpha-1,6-)-glycoprotein beta-1,2-N-acetylglucosaminyltransf | Yes |
| 55320 | MIS18BP1 | 14q21.1 | MIS18 binding protein 1 | Yes |
| 4288 | MKI67 | 10q26.2 | marker of proliferation Ki-67 | Yes |
| 84057 | MND1 | 4q31.3 | meiotic nuclear divisions 1 homolog (S. cerevisiae) | Yes |
| 219972 | MPEG1 | 11q12.1 | macrophage expressed 1 | Yes |
| 4436 | MSH2 | 2p21 | mutS homolog 2 | Yes |
| 4439 | MSH5 | 6p21.3 | mutS homolog 5 | Yes |

| | | | | |
|---|---|---|---|---|
| 339287 | MSL1 | 17q21.1 | male-specific lethal 1 homolog (Drosophila) | Yes |
| 4609 | MYC | 8q24 | v-myc avian myelocytomatosis viral oncogene homolog | Yes |
| 440145 | MZT1 | 13q22.1 | mitotic spindle organizing protein 1 | Yes |
| 4678 | NASP | 1p34.1 | nuclear autoantigenic sperm protein (histone-binding) | Yes |
| 23397 | NCAPH | 2q11.2 | non-SMC condensin I complex, subunit H | Yes |
| 8202 | NCOA3 | 20q12 | nuclear receptor coactivator 3 | Yes |
| 10403 | NDC80 | 18p11.31 | NDC80 kinetochore complex component | Yes |
| 54820 | NDE1 | 16p13.11 | nudE neurodevelopment protein 1 | Yes |
| 55247 | NEIL3 | 4q34 | nei endonuclease VIII-like 3 (E. coli) | Yes |
| 4751 | NEK2 | 1q32.3 | NIMA-related kinase 2 | Yes |
| 4780 | NFE2L2 | 2q31 | nuclear factor, erythroid 2-like 2 | Yes |
| 55655 | NLRP2 | 19q13.42 | NLR family, pyrin domain containing 2 | Yes |
| 4837 | NNMT | 11q23.1 | nicotinamide N-methyltransferase | Yes |
| 2063 | NR2F6 | 19p13.11 | nuclear receptor subfamily 2, group F, member 6 | Yes |
| 83540 | NUF2 | 1q23.3 | NUF2, NDC80 kinetochore complex component | Yes |
| 129401 | NUP35 | 2q32 | nucleoporin 35kDa | Yes |
| 51203 | NUSAP1 | 15q14 | nucleolar and spindle associated protein 1 | Yes |
| 10482 | NXF1 | 11q12-q13 | nuclear RNA export factor 1 | Yes |
| 4973 | OLR1 | 12p13.1-p12.3 | oxidized low density lipoprotein (lectin-like) receptor 1 | Yes |
| 23596 | OPN3 | 1q43 | opsin 3 | Yes |
| 4998 | ORC1 | 1p32 | origin recognition complex, subunit 1 | Yes |
| 4999 | ORC2 | 2q33 | origin recognition complex, subunit 2 | Yes |
| 23595 | ORC3 | 6q | origin recognition complex, subunit 3 | Yes |
| 5000 | ORC4 | 2q22-q23 | origin recognition complex, subunit 4 | Yes |
| 5001 | ORC5 | 7q22.1 | origin recognition complex, subunit 5 | Yes |
| 23594 | ORC6 | 16q12 | origin recognition complex, subunit 6 | Yes |
| 124222 | PAQR4 | 16p13 | progestin and adipoQ receptor family member IV | Yes |
| 55872 | PBK | 8p21.2 | PDZ binding kinase | Yes |
| 5093 | PCBP1 | 2p13-p12 | poly(rC) binding protein 1 | Yes |
| 51585 | PCF11 | 11q13 | PCF11 cleavage and polyadenylation factor subunit | Yes |
| 5111 | PCNA | 20p13-p12.3 | proliferating cell nuclear antigen | Yes |
| 80119 | PIF1 | 15q22.1 | PIF1 5'-to-3' DNA helicase | Yes |
| 9088 | PKMYT1 | 16p13.3 | protein kinase, membrane associated tyrosine/threonine 1 | Yes |
| 5324 | PLAG1 | 8q12 | pleiomorphic adenoma gene 1 | Yes |
| 55344 | PLCXD1 | Xp22.33 and Yp11.32 | hosphatidylinositol-specific phospholipase C, X domain containing | Yes |
| 153478 | PLEKHG4B | 5p15.33 | n homology domain containing, family G (with RhoGef domain) me | Yes |
| 5347 | PLK1 | 16p | polo-like kinase 1 | Yes |
| 55629 | PNRC2 | 1p36.11 | proline-rich nuclear receptor coactivator 2 | Yes |
| 10714 | POLD3 | 11q14 | polymerase (DNA-directed), delta 3, accessory subunit | Yes |
| 10721 | POLQ | 3q13.3 | polymerase (DNA directed), theta | Yes |
| 5514 | PPP1R10 | 6p21.3 | protein phosphatase 1, regulatory subunit 10 | Yes |
| 9055 | PRC1 | 15q26.1 | protein regulator of cytokinesis 1 | Yes |
| 5591 | PRKDC | 8q11 | protein kinase, DNA-activated, catalytic polypeptide | Yes |
| 55771 | PRR11 | 17q23.2 | proline rich 11 | Yes |
| 84262 | PSMG3 | 7p22.3 | proteasome (prosome, macropain) assembly chaperone 3 | Yes |
| 84722 | PSRC1 | 1p13.3 | proline/serine-rich coiled-coil 1 | Yes |
| 9232 | PTTG1 | 5q35.1 | pituitary tumor-transforming 1 | Yes |
| 10744 | PTTG2 | 4p14 | pituitary tumor-transforming 2 | Yes |
| 11137 | PWP1 | 12q23.3 | PWP1 homolog (S. cerevisiae) | Yes |
| 5885 | RAD21 | 8q24.11 | RAD21 homolog (S. pombe) | Yes |
| 10635 | RAD51AP1 | 12p13.2-p13.1 | RAD51 associated protein 1 | Yes |
| 5889 | RAD51C | 17q25.1 | RAD51 paralog C | Yes |
| 8438 | RAD54L | 1p32 | RAD54-like (S. cerevisiae) | Yes |
| 5901 | RAN | 12q24.33 | RAN, member RAS oncogene family | Yes |
| 5905 | RANGAP1 | 22q13 | Ran GTPase activating protein 1 | Yes |
| 5925 | RB1 | 13q14.2 | retinoblastoma 1 | Yes |
| 5932 | RBBP8 | 18q11.2 | retinoblastoma binding protein 8 | Yes |
| 5933 | RBL1 | 20q11.23 | retinoblastoma-like 1 | Yes |
| 5934 | RBL2 | 16q12.2 | retinoblastoma-like 2 | Yes |
| 9978 | RBX1 | 22q13.2 | ring-box 1, E3 ubiquitin protein ligase | Yes |
| 1827 | RCAN1 | 21q22.1-q22.2 | regulator of calcineurin 1 | Yes |
| 5982 | RFC2 | 7q11.23 | replication factor C (activator 1) 2, 40kDa | Yes |
| 5984 | RFC4 | 3q27 | replication factor C (activator 1) 4, 37kDa | Yes |
| 5998 | RGS3 | 9q32 | regulator of G-protein signaling 3 | Yes |
| 83695 | RHNO1 | 12p13.33 | RAD9-HUS1-RAD1 interacting nuclear orphan 1 | Yes |
| 22836 | RHOBTB3 | 5q15 | Rho-related BTB domain containing 3 | Yes |
| 114822 | RHPN1 | 8q24.3 | rhophilin, Rho GTPase binding protein 1 | Yes |
| 6167 | RPL37 | 5p13.1 | ribosomal protein L37 | Yes |
| 6241 | RRM2 | 2p25-p24 | ribonucleotide reductase M2 | Yes |
| 8568 | RRP1 | 21q22.3 | ribosomal RNA processing 1 | Yes |
| 57035 | RSRP1 | 1p36.13-p35.1 | arginine/serine-rich protein 1 | Yes |
| 861 | RUNX1 | 21q22.3 | runt-related transcription factor 1 | Yes |

| | | | | |
|---|---|---|---|---|
| 8819 | SAP30 | 4q34.1 | Sin3A-associated protein, 30kDa | Yes |
| 89958 | SAPCD2 | 9q34.3 | suppressor APC domain containing 2 | Yes |
| 57410 | SCYL1 | 11q11-q12 | SCY1-like 1 (S. cerevisiae) | Yes |
| 22929 | SEPHS1 | 10p14 | selenophosphate synthetase 1 | Yes |
| 57190 | SEPN1 | 1p36.13 | selenoprotein N, 1 | No |
| 6318 | SERPINB4 | 18q21.33 | serpin peptidase inhibitor, clade B (ovalbumin), member 4 | Yes |
| 2810 | SFN | 1p36.11 | stratifin | Yes |
| 6421 | SFPQ | 1p34.3 | splicing factor proline/glutamine-rich | Yes |
| 6444 | SGCD | 5q33-q34 | sarcoglycan, delta (35kDa dystrophin-associated glycoprotein) | Yes |
| 151246 | SGOL2 | 2q33.2 | shugoshin-like 2 (S. pombe) | No |
| 6456 | SH3GL2 | 9p22 | SH3-domain GRB2-like 2 | Yes |
| 79801 | SHCBP1 | 16q11 | SHC SH2-domain binding protein 1 | Yes |
| 221150 | SKA3 | 13q11 | spindle and kinetochore associated complex subunit 3 | Yes |
| 6500 | SKP1 | 5q31 | S-phase kinase-associated protein 1 | Yes |
| 6502 | SKP2 | 5p13 | S-phase kinase-associated protein 2, E3 ubiquitin protein ligase | Yes |
| 7884 | SLBP | 4p16.3 | stem-loop binding protein | Yes |
| 10246 | SLC17A2 | 6p22.2 | solute carrier family 17, member 2 | Yes |
| 6507 | SLC1A3 | 5p13 | ıte carrier family 1 (glial high affinity glutamate transporter), membe | Yes |
| 123096 | SLC25A29 | 14q32.2 | carrier family 25 (mitochondrial carnitine/acylcarnitine carrier), mem | Yes |
| 57181 | SLC39A10 | 2q33.1 | solute carrier family 39 (zinc transporter), member 10 | Yes |
| 4087 | SMAD2 | 18q21 | SMAD family member 2 | Yes |
| 4088 | SMAD3 | 15q21-q22 | SMAD family member 3 | Yes |
| 4089 | SMAD4 | 18q21.1 | SMAD family member 4 | Yes |
| 8243 | SMC1A | Xp11.22-p11.21 | structural maintenance of chromosomes 1A | Yes |
| 27127 | SMC1B | 22q13 | structural maintenance of chromosomes 1B | Yes |
| 9126 | SMC3 | 10q25 | structural maintenance of chromosomes 3 | Yes |
| 10051 | SMC4 | 3q26.1 | structural maintenance of chromosomes 4 | Yes |
| 6525 | SMTN | 22q12 | smoothelin | Yes |
| 8470 | SORBS2 | 4q35.1 | sorbin and SH3 domain containing 2 | Yes |
| 10615 | SPAG5 | 17q11.2 | sperm associated antigen 5 | Yes |
| 54908 | SPDL1 | 5q35.1 | spindle apparatus coiled-coil protein 1 | Yes |
| 6715 | SRD5A1 | 5p15.31 | reductase, alpha polypeptide 1 (3-oxo-5 alpha-steroid delta 4-dehydr | Yes |
| 140890 | SREK1 | 5q11.2-q12.1 | splicing regulatory glutamine/lysine-rich protein 1 | Yes |
| 6428 | SRSF3 | 6p21 | serine/arginine-rich splicing factor 3 | Yes |
| 10274 | STAG1 | 3q22.2-q22.3 | stromal antigen 1 | Yes |
| 10735 | STAG2 | Xq25 | stromal antigen 2 | Yes |
| 6772 | STAT1 | 2q32.2-q32.3 | signal transducer and activator of transcription 1, 91kDa | Yes |
| 6777 | STAT5B | 17q11.2 | signal transducer and activator of transcription 5B | Yes |
| 6491 | STIL | 1p32 | SCL/TAL1 interrupting locus | Yes |
| 9262 | STK17B | 2q33.1 | serine/threonine kinase 17b | Yes |
| 8801 | SUCLG2 | 3p14.3 | succinate-CoA ligase, GDP-forming, beta subunit | Yes |
| 10460 | TACC3 | 4p16.3 | transforming, acidic coiled-coil containing protein 3 | Yes |
| 11138 | TBC1D8 | 2q12.1 | TBC1 domain family, member 8 (with GRAM domain) | Yes |
| 6924 | TCEB3 | 1p36.1 | ıscription elongation factor B (SIII), polypeptide 3 (110kDa, elongin | No |
| 7027 | TFDP1 | 13q34 | transcription factor Dp-1 | Yes |
| 7029 | TFDP2 | 3q23 | transcription factor Dp-2 (E2F dimerization partner 2) | Yes |
| 7040 | TGFB1 | 19q13.1 | transforming growth factor, beta 1 | Yes |
| 7042 | TGFB2 | 1q41 | transforming growth factor, beta 2 | Yes |
| 7043 | TGFB3 | 14q24 | transforming growth factor, beta 3 | Yes |
| 7076 | TIMP1 | Xp11.3-p11.23 | TIMP metallopeptidase inhibitor 1 | Yes |
| 51014 | TMED7 | 5q22.3 | transmembrane emp24 protein transport domain containing 7 | Yes |
| 7153 | TOP2A | 17q21-q22 | topoisomerase (DNA) II alpha 170kDa | Yes |
| 11073 | TOPBP1 | 3q22.1 | topoisomerase (DNA) II binding protein 1 | Yes |
| 7157 | TP53 | 17p13.1 | tumor protein p53 | Yes |
| 22974 | TPX2 | 20q11.2 | TPX2, microtubule-associated | Yes |
| 9319 | TRIP13 | 5p15 | thyroid hormone receptor interactor 13 | Yes |
| 27037 | TRMT2A | 22q11.21 | tRNA methyltransferase 2 homolog A (S. cerevisiae) | Yes |
| 10024 | TROAP | 12q13.12 | trophinin associated protein | Yes |
| 8848 | TSC22D1 | 13q14 | TSC22 domain family, member 1 | Yes |
| 55020 | TTC38 | 22q13 | tetratricopeptide repeat domain 38 | Yes |
| 7272 | TTK | 6q14.1 | TTK protein kinase | Yes |
| 7846 | TUBA1A | 12q13.12 | tubulin, alpha 1a | Yes |
| 113457 | TUBA3D | 2q21.1 | tubulin, alpha 3d | Yes |
| 7277 | TUBA4A | 2q36.1 | tubulin, alpha 4a | Yes |
| 203068 | TUBB | 6p21.33 | tubulin, beta class I | Yes |
| 7280 | TUBB2A | 6p25.2 | tubulin, beta 2A class IIa | Yes |
| 10383 | TUBB4B | 9q34.3 | tubulin, beta 4B class IVb | Yes |
| 51174 | TUBD1 | 17q23.1 | tubulin, delta 1 | Yes |
| 11065 | UBE2C | 20q13.12 | ubiquitin-conjugating enzyme E2C | Yes |
| 7323 | UBE2D3 | 4q24 | ubiquitin-conjugating enzyme E2D 3 | Yes |
| 140739 | UBE2F | 2q37.3 | ubiquitin-conjugating enzyme E2F (putative) | Yes |
| 27338 | UBE2S | 19q13.43 | ubiquitin-conjugating enzyme E2S | Yes |

| | | | | |
|---|---|---|---|---|
| 29089 | UBE2T | 1q32.1 | ubiquitin-conjugating enzyme E2T | Yes |
| 51366 | UBR5 | 8q22 | ubiquitin protein ligase E3 component n-recognin 5 | Yes |
| 55148 | UBR7 | 14q32.12 | ubiquitin protein ligase E3 component n-recognin 7 (putative) | Yes |
| 7374 | UNG | 12q23-q24.1 | uracil-DNA glycosylase | Yes |
| 79650 | USB1 | 16q13 | U6 snRNA biogenesis 1 | Yes |
| 7398 | USP1 | 1p31.3 | ubiquitin specific peptidase 1 | Yes |
| 81839 | VANGL1 | 1p13.1 | VANGL planar cell polarity protein 1 | Yes |
| 84313 | VPS25 | 17q21.31 | vacuolar protein sorting 25 homolog (S. cerevisiae) | Yes |
| 79968 | WDR76 | 15q15.3 | WD repeat domain 76 | Yes |
| 197335 | WDR90 | 16p13.3 | WD repeat domain 90 | Yes |
| 7465 | WEE1 | 11p15.4 | WEE1 G2 checkpoint kinase | Yes |
| 494551 | WEE2 | 7q32 | WEE1 homolog 2 (S. pombe) | Yes |
| 8840 | WISP1 | 8q24.22 | WNT1 inducible signaling pathway protein 1 | No |
| 26118 | WSB1 | 17q11.2 | WD repeat and SOCS box containing 1 | Yes |
| 7494 | XBP1 | 22q12.1 | X-box binding protein 1 | Yes |
| 64328 | XPO4 | 13q11 | exportin 4 | Yes |
| 9213 | XPR1 | 1q25.1 | xenotropic and polytropic retrovirus receptor 1 | Yes |
| 8089 | YEATS4 | 12q13-q15 | YEATS domain containing 4 | Yes |
| 7529 | YWHAB | 20q13.1 | ıe 3-monooxygenase/tryptophan 5-monooxygenase activation protei | Yes |
| 7531 | YWHAE | 17p13.3 | 3-monooxygenase/tryptophan 5-monooxygenase activation protein, | Yes |
| 7532 | YWHAG | 7q11.23 | 3-monooxygenase/tryptophan 5-monooxygenase activation protein, | Yes |
| 7533 | YWHAH | 22q12.1-q13.1 | ne 3-monooxygenase/tryptophan 5-monooxygenase activation prote | Yes |
| 10971 | YWHAQ | 2p25.2-p25.1 | e 3-monooxygenase/tryptophan 5-monooxygenase activation protei | Yes |
| 7534 | YWHAZ | 8q22.3 | ıe 3-monooxygenase/tryptophan 5-monooxygenase activation protei | Yes |
| 7709 | ZBTB17 | 1p36.13 | zinc finger and BTB domain containing 17 | Yes |
| 23144 | ZC3H3 | 8q24.3 | zinc finger CCCH-type containing 3 | Yes |
| 51530 | ZC3HC1 | 7q32.2 | zinc finger, C3HC-type containing 1 | Yes |
| 23174 | ZCCHC14 | 16q24.2 | zinc finger, CCHC domain containing 14 | Yes |
| 57209 | ZNF248 | 10p11.21 | zinc finger protein 248 | Yes |
| 195828 | ZNF367 | 9q22 | zinc finger protein 367 | Yes |
| 84330 | ZNF414 | 19p13.2 | zinc finger protein 414 | Yes |
| 25925 | ZNF521 | 18q11.2 | zinc finger protein 521 | Yes |
| 11055 | ZPBP | 7p14.3 | zona pellucida binding protein | Yes |
| 9406 | ZRANB2 | 1p31 | zinc finger, RAN-binding domain containing 2 | Yes |
| 11130 | ZWINT | 10q21-q22 | ZW10 interacting kinetochore protein | Yes |

Genes are ordered alphabbetically
