## Supplemental Table 2. for "Reclassifying Cancer: Defining tumour cell cycle activity in terms of its tissue of origin in over 13,000 samples"

Tumour purity level among different cancer types

| **Tissue of origin** | **Cancer type** | **Avg. purity** |
|---|---|---|
| Adrenal Gland | ACC | 0.83 |
| Bladder | BLCA | 0.62 |
| Brain | LGG | 0.72 |
| Brain | GBM | 0.76 |
| Breast | BRCA | 0.61 |
| Cervix | CESC | 0.66 |
| Colon | COAD | 0.64 |
| Endometrium | UCEC | 0.73 |
| Esophagus | ESCA | 0.63 |
| Head and Neck | HNSC | 0.52 |
| Kidney | KIRC | 0.58 |
| Kidney | KIRP | 0.74 |
| Kidney | KICH | 0.82 |
| Liver | LIHC | 0.7 |
| Lung | LUAD | 0.47 |
| Lung | LUSC | 0.52 |
| Ovary | OV | 0.78 |
| Pancreas | PAAD | 0.55 |
| Prostate | PRAD | 0.62 |
| Skin | SKCM | 0.68 |
| Stomach | STAD | 0.53 |
| Thyroid Gland | THCA | 0.72 |
| Uterus | UCS | 0.82 |

Avg. purity: Average tumour purity, ACC: Adrenocortical carcinoma, BLCA: Bladder Urothelial Carcinoma, BRCA: Breast invasive carcinoma, CESC: Cervical squamous cell carcinoma and endocervical adenocarcinoma, COAD: Colon adenocarcinoma, ESCA: Esophageal carcinoma, GBM: Glioblastoma multiforme, HNSC: Head and Neck squamous cell carcinoma, KICH: Kidney Chromophobe, KIRC: Kidney renal clear cell carcinoma, KIRP: Kidney renal papillary cell carcinoma, LGG: Brain Lower Grade Glioma, LIHC: Liver hepatocellular carcinoma, LUAD: Lung adenocarcinoma, LUSC: Lung squamous cell carcinoma, OV: Ovarian serous cystadenocarcinoma, PAAD: Pancreatic adenocarcinoma, PRAD: Prostate adenocarcinoma, SKCM: Skin Cutaneous Melanoma, STAD: Stomach adenocarcinoma, THCA: Thyroid carcinoma, THYM: Thymoma, UCS: Uterine Carcinosarcoma
