## Supplemental Table 3. for "Reclassifying Cancer: Defining tumour cell cycle activity in terms of its tissue of origin in over 13,000 samples"

Supplemental table 3.
Number of outliers according to PCA

| Tissue of origin | Variable genes | | CCS | |
|---|---|---|---|---|
| | Normal | Tumour | Normal | Tumour |
| Adrenal Gland | 0 | 0 | 1 | 0 |
| Brain | 0 | 0 | 6 | 0 |
| Colon | 0 | 0 | 4 | 0 |
| Esophagus | 0 | 0 | 4 | 0 |
| Kidney | 0 | 0 | 1 | 0 |
| Lung | 0 | 0 | 1 | 1 |
| Ovary | 0 | 0 | 0 | 1 |
| Pancreas | 0 | 0 | 2 | 0 |
| Stomach | 0 | 0 | 2 | 0 |
| Thyroid Gland | 0 | 0 | 1 | 0 |
| Testis | 156 | 163 | 0 | 0 |
| Total | 156 | 163 | 22 | 2 |
| | 319 | | 24 | |
